# Different resource partitioning explains plant species richness patterns in tropical alpine ecosystems

**DOI:** 10.1101/2024.06.24.599547

**Authors:** Martha Kandziora, Diana L. A. Vásquez, Christian Brochmann, Abel Gizaw, Lovisa Gustafsson, Desalegn Chala, Mercè Galbany-Casals, Filip Kolář, Petr Sklenář, Nicolai M. Nürk, Roswitha Schmickl

## Abstract

Species co-existence based on resource partitioning modulates biodiversity patterns across latitudes and altitudes. Resource partitioning can occur via specialisation or separation in the geographic range or niche. Here, we compare two tropical alpine ecosystems with similar climates to test for geographic range and climatic niche partitioning strategies in explaining species richness difference. We compare the species-rich tropical alpine ecosystem in the South American Andes with the more species-poor one in the eastern African mountains. We combine phylogenomic data for three locally diversified plant lineages in each region with occurrence records and estimate climatic niche and geographic range metrics (size and overlap). We found that the Andean species have overall larger niches than the African species, thus smaller niches indicating specialisation is not the explanation for the higher species richness in the Andes. Instead, for species with overlapping geographic ranges, we found that the Andean species tend to show less niche overlap than the African species, indicating more effective niche separation. Taken together, we propose that different degrees of niche separation in geographically overlapping species, and hence, a different pattern of resource partitioning, explain the differences in species richness between the two tropical alpine ecosystems.

## Introduction

Regions with similar physical and environmental features are expected to harbour similar species numbers per area^1^. The concept of niche, describing conditions for species survival, is crucial for understanding species distribution in space and time^2^. Geodiversity and climate accounts for 50-70% of the variation in plant species richness^3^. Especially mountains, with their large variation in geodiversity and climate, harbour an exceptionally high concentration of species relative to their surface area^3–7^.

Species inhabiting the same general area, depend on resource partitioning to minimise competition, and this promotes species richness^8–10^. Resource partitioning can arise via range and niche partitioning, i.e. via niche specialisation, small range sizes^11,12^, niche separation, and/or range separation. Alternatively, species can minimise competition through partial resource partitioning, which can occur via different degrees of niche packing^13–15^. Niche preferences tend to be retained during speciation^16^, but when closely related species do show divergent niches this may permit a higher number of species to coexist. The concept of resource partitioning increasing biodiversity is well established, but it remains unclear whether differences in niche specialisation, range size, niche separation, range separation, or niche evolution can explain why regions with similar physical and environmental features may differ markedly in species richness, even when accounting for area and time.

Tropical alpine ecosystems present a unique opportunity to investigate species richness differences, as they have similar climatic conditions^17^ and are of similar age^18,19^. Furthermore, these ecosystems are cold continental islands surrounded by warm tropical lowlands, and they are found on different continents. Hence, most of the flora comprises lineages with non-tropical origins^20^, as species require certain adaptations to enable colonisation^20,21^, the majority of immigrants originated from non-tropical sources^20^. As such, species richness differences among tropical alpine ecosystems^22^ should be independent of those among tropical regions *per se*. Nevertheless, despite their similarities, tropical alpine ecosystems exhibit conspicuous disparities in plant species richness, habitat size and connectivity^17,20,23,24^, with the latter two likely having an effect on plant species richness.

Here, we compare two tropical alpine ecosystems to test for patterns in niche and range partitioning potentially explaining the plant species richness differences among similar ecosystems (Fig. 1): the large, well-connected, and species-rich Páramo region in the northern Andes in South America, and the smaller, highly fragmented and relatively species-poor Afroalpine region in the eastern African mountains. Notably, species richness is higher in the Páramo, even when correcting for area^25,26^, and when calculating species densities within plots^27^ and species accumulation curves^28^. We constructed and analysed time-scaled phylogenies of three prominent and locally diversified plant lineages from each ecosystem and compared niche and range metrics. Using phylogenetic comparative methods, we tested whether species richness differences are associated with niche specialisation or small ranges (measured here as niche and range size; hypothesis 1), with range separation or niche packing (measured as range overlap and as niche overlap of range-overlapping species; hypothesis 2), or with niche evolution (measured as distances between niche axis optima compared to genetic distances; hypothesis 3).

**Figure 1.**
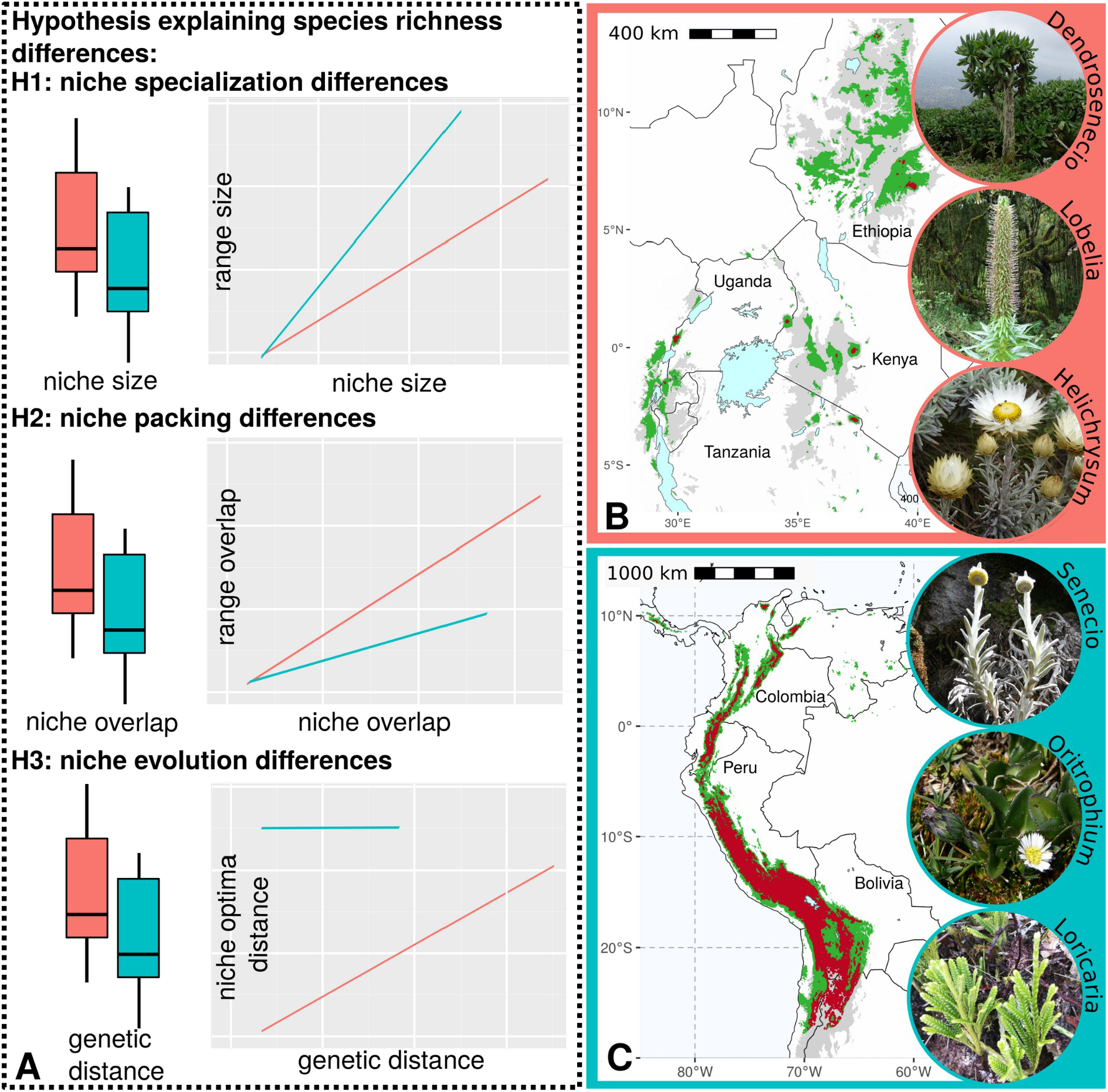
Conceptual figure showing niche and range hypotheses explaining species richness differences between the two tropical alpine ecosystems. (A) Potential patterns of niche and range metrics explaining species richness differences in the species-poor Afroalpine (red) and the species-rich Páramo (blue). We hypothesise that species richness is higher in the Páramo because (H1) niche specialisation is higher (smaller niche sizes; in addition, because of its larger area, range size will increase more steeply with niche size); (H2) higher species turnover and hence lower levels of niche packing (niche and range overlap is lower); or (H3) higher niche evolution drives species evolution and permits more species to coexist (smaller genetic distances and larger niche optima distances between species). Maps highlighting the tropical alpine ecosystems, based on a climatic delimitation^17^, of the eastern African mountains (Afroalpine; B) and the South American Andes (Páramo; C). Tropical alpine conditions are shown in red, green delimits temperate conditions based on climate. Grey colours designate areas above 1500 m above sea level , areas below are white. Circular insets show the characteristic, locally diversified plant lineages analysed here. Images provided by Martha Kandziora for the Afroalpine and by Petr Sklenář for the Páramo species.

## Results

### Phylogenetic reconstruction of three lineages in each tropical alpine ecosystem

We inferred phylogenies based on target enrichment sequence data. In South America we selected the genera *Oritrophium* (Kunth) Cuatrec. (12 of 18 accepted species), *Loricaria* Wedd. (13 of 19 accepted species^29^), and a clade of *Senecio* L. (ser. *Culcitium*; 36 species out of ca. 41 accepted species) as Páramo representatives. In Africa we selected the genus *Dendrosenecio* (Hauman ex Hedberg) B. Nord. (including all 11 accepted species^30^), a tropical alpine clade of *Helichrysum* Mill. (including all 10 accepted species^31^), and a tropical alpine clade of giant lobelias (*Lobelia* L., including all 13 accepted species^32^) as representatives of the Afroalpine. Five of the six lineages belong to Asteraceae, the most species-rich family in tropical alpine ecosystems in general^26,33^. For these five lineages phylogenies existed already^29–31,34,35^. Nevertheless, we recalculated three of these five phylogenies in order to confirm monophyly of species and to add newly sampled taxa while removing samples showing gene flow (Supplementary Note S1; Supplementary Figs. S1- S3). For the giant lobelia clade (Campanulaceae), we generated a new phylogeny (Supplementary Fig. S4; Supplementary Note S1 and S2). Using a penalised likelihood approach^36,37^, we estimated that all lineages diversified from the late Miocene to the early Pliocene, which is in accordance with previous works^19,31,38–40^ (crown ages range from 7.11 to 3.89 million years ago; Supplementary Figs. S5-S10; Supplementary Table S1). These estimates demonstrate that our analyses of niche and range metrics compare lineages of similar evolutionary ages.

### Niche packing, not niche specialisation, drives species richness differences

For the species-wide multi-dimensional climatic niche metrics, we collected occurrence data from herbarium specimens supplemented with data from GBIF (6 to 184 occurrences per species, with a mean of 26 to 49 species occurrence points per lineage). After excluding species represented by less than six occurrences, we calculated niche and range information for 32 African (1025 occurrences in total) and 52 Andean species (1917 occurrences in total; Supplementary Table S2). We excluded 24 rare Andean species because of insufficient occurrence information (*Loricaria*: six species; *Oritrophium*: six species; *Senecio*: 12 species) and one Afroalpine species because it only occurs outside the tropical montane regions.

We calculated niche size, niche optima, niche overlap, and new niche space (niche expansion plus unfilling divided by two) based on ordination of present-day bioclimatic variables from the CHELSA^41^ global climate data grids. We selected eight climatic variables based on the Variation Inflation Factor (VIF) using a stepwise procedure to reduce correlation among variables (VIF < 9; Supplementary Table S3; Supplementary Fig. S11). The niche metrics of each species were inferred from the first two axes of a principal component analysis (PCAenv following ^42^), which accounted for ∼80% of the total variation (PC1 59.0%, PC2 20.4%; Supplementary Fig. S12). The bioclimatic variables that primarily defined PC1 comprised precipitation- and seasonality-related variables, while PC2 mainly depicted temperature-related variables. Geographic range metrics were calculated based on the minimum extent encompassing all occurrences per species and removing areas with tropical climate within this minimum extent to constrain ranges to temperate climate^17^.

Niche size, a proxy for niche specialisation, was significantly larger in the Páramo species than in the Afroalpine species (p = 0.012; Fig. 2A; Supplementary Table S4). This was most pronounced along PC1, designating the precipitation and seasonality of the niche. This niche axis in particular revealed small niche sizes for the Afroalpine *Dendrosenecio* species, but large sizes for the Andean *Senecio* species (Supplementary Fig. S13; Supplementary Table S5). The Páramo species also had overall larger range sizes than the Afroalpine species (p = 1.51e-05; Fig. 2B; Supplementary Table S4). To test for significance in correlation between range size and niche size, we used phylogenetic generalised least squares (PGLS) models, which revealed (1) positive correlations for each lineage (p < 0.05), and (2) that niche size was mainly explained by the precipitation- and seasonality-related PC1 axis (Fig. 2C; Supplementary Table S6). The adjusted R^2^, designating the proportion of the variation in the dependent variable that is predictable from the independent variable, explained 41-77% of the variance per lineage (Supplementary Table S6).

**Figure 2.**
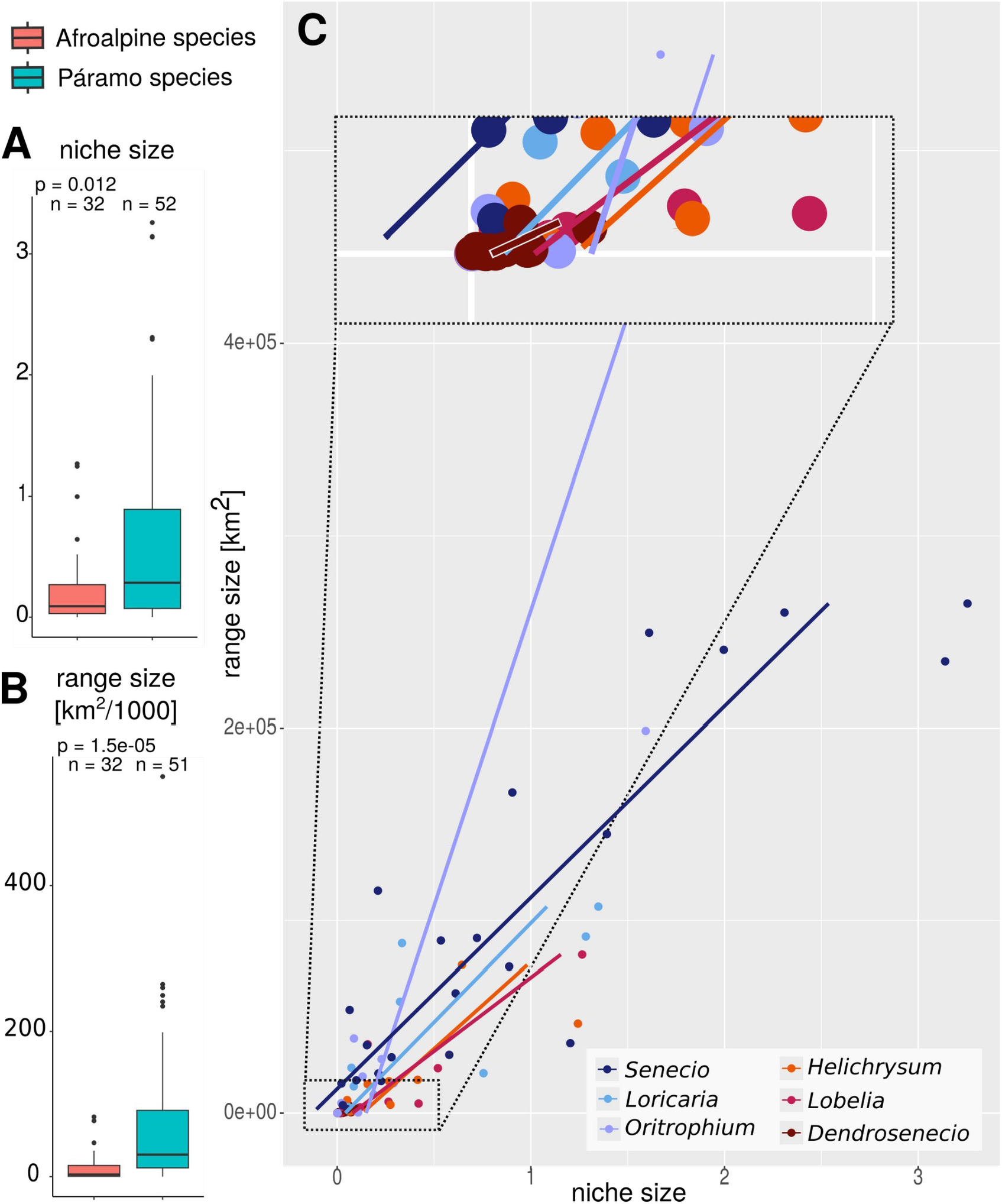
Climatic niche and range size of the Afroalpine and Páramo species. (A) Boxplots of niche size and (B) range size summarised for all species in each region. Boxplots show the median (centre line) and the first and third quartiles, while whiskers mark the 5th and 95th percentiles, values beyond these bounds are outliers. The p-values were calculated using a Wilcoxon test. Abbreviation: n - number of data points. (C) Phylogenetic generalised least squares of range size and niche size of each lineage; all were significant (Supplementary Table S6). Enlarged inset to increase visibility of the small niche sizes. Contrary to expectations of hypothesis 1, the niche size of Páramo species was larger compared to Afroalpine species, indicating less climatic specialisation.

We found that the degree of niche overlap and range overlap were similar in the Páramo and Afroalpine species (Fig. 3A, 3B; Supplementary Table S4). Niche overlap and range overlap appeared spatially autocorrelated, somewhat more strongly for the Afroalpine species (Fig. 3C), with the highest degree of overlap in the Afroalpine *Helichrysum* (Supplementary Fig. E9; Supplementary Table S5). We measured the degree of niche packing^14,43^ as niche overlap on the subset of species with overlapping geographic ranges (set as >10% range overlap to exclude marginally range-overlapping species). For the subset of species with non-overlapping ranges, we found similar degrees of niche overlap in the Páramo and the Afroalpine (p = 0.116; Fig. 3C, 3D; Supplementary Table S4). In contrast, in range-overlapping species, the Páramo species had significantly lower niche overlap, i.e. lower degree of niche packing, than the Afroalpine species (p = 0.014; Fig. 3C, 3D; Supplementary Table S4).

**Figure 3.**
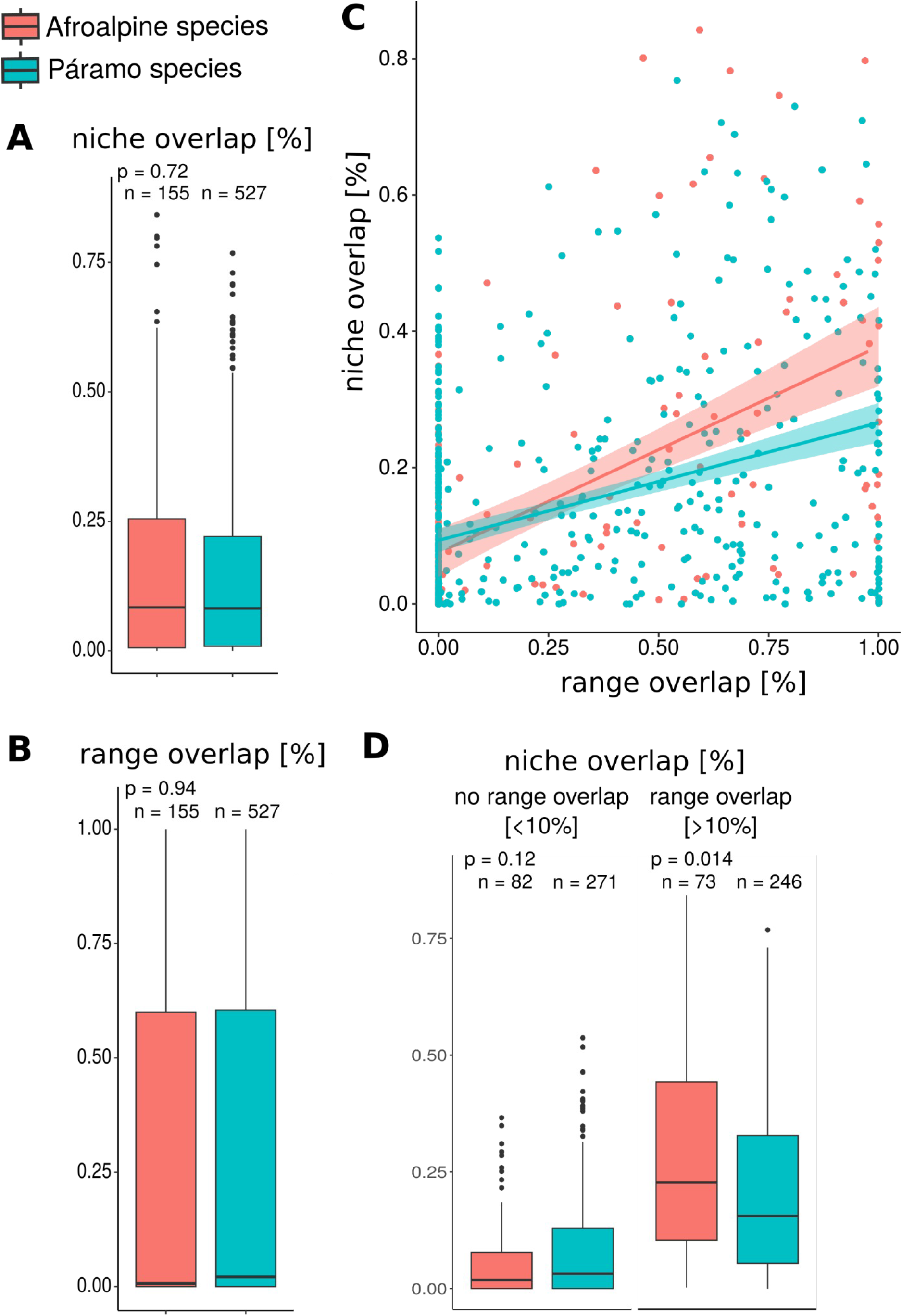
Climatic niche and range overlap of the Afroalpine and Páramo plant lineages. (A) Boxplots of niche overlap and (B) range overlap calculated for all species in each lineage and then summarised per region. (C) Linear regression of range and niche overlap. (D) Boxplots of niche overlap calculated for the subsets of species with non-overlapping and overlapping ranges. Boxplots show the median (centre line) and the first and third quartiles, while whiskers mark the 5th and 95th percentiles, values beyond these bounds are outliers. The p-values were calculated using a Wilcoxon test. Abbreviation: n - number of data points per subset. Results agree with expectations of hypothesis 2, there was a lower niche overlap of range-overlapping Páramo species, indicating higher degrees of niche separation and hence lower degrees of niche packing than in Afroalpine species.

### Niche evolution differences between the two tropical alpine ecosystems

We examined niche evolution, approximated by niche distances regressed to genetic distances, in all 32 Afroalpine species and in 43 Páramo species for which occurrence and phylogenetic data were available. The genetic distance, measured as cophenetic distance in million years between species within a lineage, was smaller in the Páramo lineages than in the Afroalpine lineages (p = 0.00025), as expected due to larger numbers of species per lineage in the Páramo, whereas the crown ages of lineages were similar across regions (Supplementary Table S4). Nevertheless, contrary to our expectations, we did not find overall differences in niche evolution between the Andean and the Afroalpine lineages (Fig. 4). In four of the six lineages, namely *Dendrosenecio* (Afroalpine) and all Páramo lineages, we found positive correlations between genetic distance and new niche space (Fig. 4A), resembling gradual niche evolution as expected under neutral evolution. In these lineages, with the exception of *Oritrophium* (Páramo), also the distance between species ranges, measured in metres, became larger with increasing genetic distance (Fig. 4B). Niche differentiation in these three lineages appeared to be related to niche optima distances of the precipitation- and seasonality-related niche axis (PC1 axis; Fig. 4C, D). Hence, in these lineages the distances between the precipitation- and seasonality-related niche optima were different between species. Whereas the median niche optima of that niche axis were similar when summarised across regions and lineages (Supplementary Table S4 and S5). In the three remaining lineages, namely *Helichrysum* and *Lobelia* (both Afroalpine) and *Oritrophium* (Páramo), we did not find any relationship between genetic distance and niche differentiation or range distance, indicating niche divergence likely resulting from ecological differentiation reducing competition.

**Figure 4.**
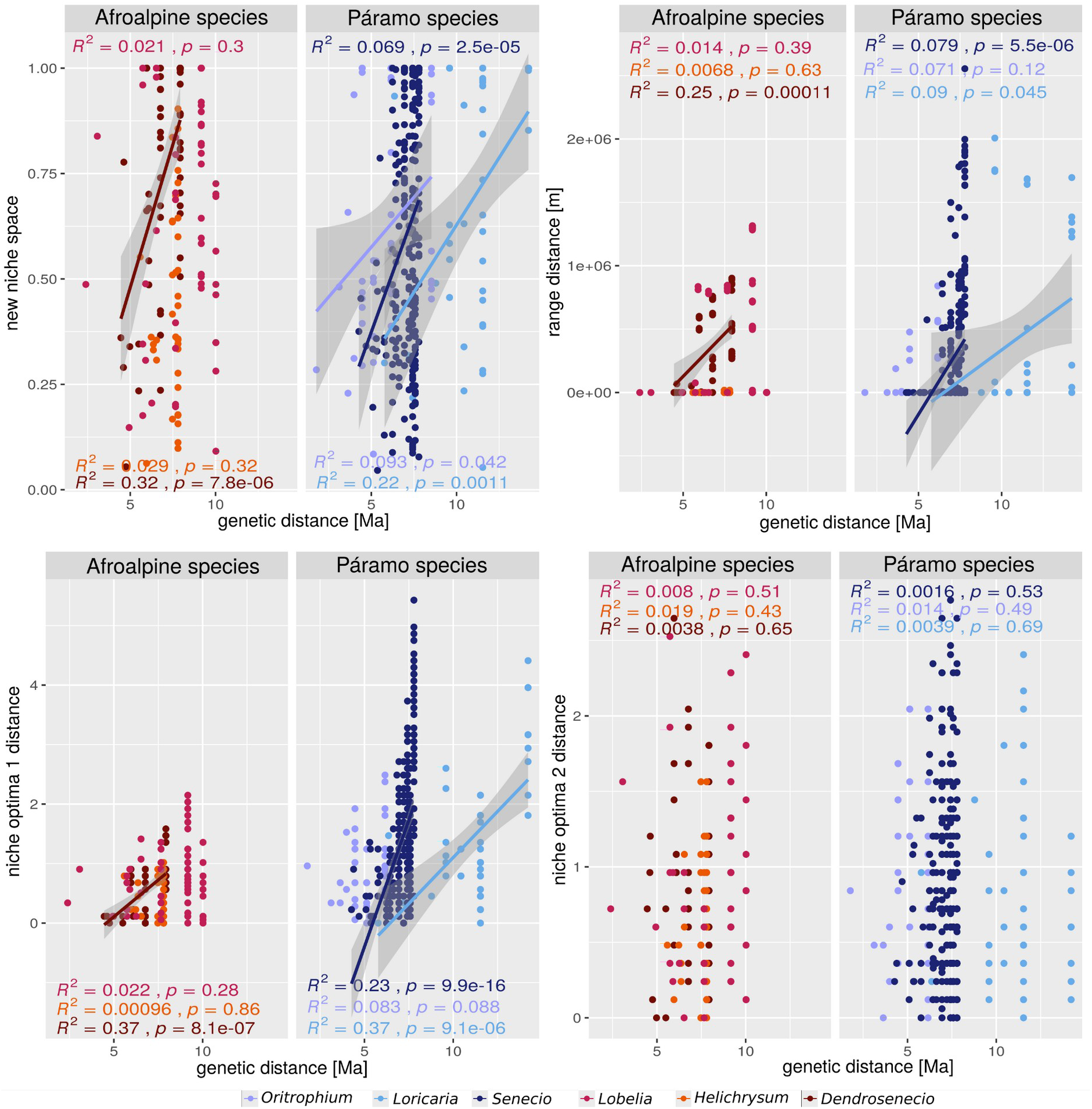
Linear regression between genetic distance and niche and range metrics, separated per lineage. Relation of genetic distance, measured as cophenetic distance in million years [Ma], to (A) new occupied niche space (niche expansion and niche unfilling divided by two), (B) the distance between species ranges measured as the closest distance between two ranges, (C) the distance between the precipitation- and seasonality-related mean niche optima (niche optima 1 distance) of two species, and (D) the distance between the temperature-related mean niche optima (niche optima 2 distance) of two species. Panels include R^2^ and p-value of linear regression (only shown if significant with a p-value <0.05 based on Mantel test) per lineage. Contrary to expectations of hypothesis 3, there was a lower correlation between genetic distances and niche and range metrics for the Afroalpine lineages than for the Páramo lineages.

## Discussion

The similarities in the evolution of the floras of the Páramo in the northern Andes and of the Afroalpine in the eastern African mountains are well documented, and prominently exemplified by the convergent evolution of giant rosette and dwarf shrub growth forms^44^. Whether the enigmatic species richness differences between the Páramo and the Afroalpine can be explained by different patterns in range and niche partitioning among species remained unexplored, mainly because of the scarcity of well-resolved phylogenies combined with limited information on species distributions. To address this gap, we compiled a comparative data set on the phylogenetic relationships, range patterns, and climatic niche preferences of species from six different plant lineages. We hypothesised that the higher species richness of the Páramo is due to increased resource partitioning, either via greater niche specialisation, lower niche packing, or higher niche evolution among the species in the studied plant lineages. Our findings reveal a complex interplay of niche and range patterns explaining species richness differences among these two tropical alpine ecosystems.

Hypothesis 1, higher species richness in the Páramo is driven by higher niche specialisation, is rejected. The examined Páramo species show larger niche and range sizes than the Afroalpine species (Fig. 2A, B). However, this difference is mainly driven by outlier clades: *Dendrosenecio* in the Afroalpine and *Senecio* in the Páramo, exhibiting small and large niche and range sizes, respectively (Supplementary Fig. E8; Supplementary Table S5). We found a positive correlation between niche and range size for species in both tropical alpine ecosystems (Fig. 2C), a pattern described for other species and ecosystems^45,46^ (but see ^47^ for a counter example). Yet, range size does not increase more steeply in Páramo than in Afroalpine species. This result is unexpected given the larger area available in the Páramo and hence, the larger area with potentially matching climate.

Small niche sizes have for example often been used to explain the latitudinal diversity gradient; stable ecosystems lead to stable populations and promote ecological specialisation over time, which can lead to smaller niche sizes and hence enables more species to coexist^11,48–50^. The negative correlation between species richness and niche size, as found in studies investigating species diversity gradients (for a review see ^11^), does not explain species richness differences between the two tropical alpine ecosystems studied here. In these two tropical alpine ecosystems niche sizes are similar (Fig. 2A, 2C), although the Páramo is more species-rich and potentially has been more climatically stable since the Last Glacial Maximum^17^. The Afroalpine region showed less climatic stability, and suitable areas are highly disconnected^17^, which might have promoted extinction of species with small niches and ranges in the Afroalpine. Hence, higher species extinction likely contributed to the overall lower species richness in the Afroalpine. In the Páramo, the higher species richness might be related to the larger available environmental space of the ecosystem (Supplementary Fig. S11). This is in line with findings by ^7^, who showed that the size of the climatic space of an ecosystem correlates with species richness. However, whether the larger climatic space present in the Páramo results from its larger extent, or whether the spatial climatic heterogeneity is higher in the Páramo than in the Afroalpine remains unexplored.

A key difference in species richness patterns in the tropical alpine ecosystems may lie in the tolerance to precipitation differences: the precipitation- and seasonality-related niche size differs significantly between the two ecosystems, with Páramo species exhibiting a broader niche (PC1 size; Supplementary Table S4). Also, overall niche size correlates with precipitation- and seasonality-variability for the studied lineages (Supplementary Table S6). In general, the tropics bear higher species richness in regions with larger precipitation variability^51,52^, implying that precipitation has a stronger effect on species richness patterns than temperature in the tropics. Hence, the higher species richness in the Páramo might arise from a higher number of species with broad climatic niches compared to the Afroalpine species, thus that there are more species that tolerate larger variation in the precipitation- and seasonality-related niche dimensions (Supplementary Table S4).

Hypothesis 2, assuming that the higher plant species richness in the Páramo is explained by higher levels of resource partitioning, here approximated by niche and range overlap patterns, is partially supported. Our findings show, first, similar levels of range distance and range overlap between species in the two regions (Fig. 3B, Supplementary Table S4), despite the vast tropical lowlands separating the scattered Afroalpine regions in contrast to the more contiguous Andean cordilleras. Second, climatic niche overlap of species is also similar in the two tropical alpine ecosystems. However, when comparing range overlapping and non-overlapping species, interesting patterns became prevalent (Fig. 3C, 3D). Niche overlap is higher in range overlapping than in non-overlapping species (Fig. 3D), reflecting spatial autocorrelation: species that occur in geographically distant areas experience more different climates than species in close geographic proximity. However, the niche overlap of range-overlapping species is lower in the Páramo than in the Afroalpine, indicating higher niche separation of Páramo species when ranges overlap (Fig. 3D).

Hypothesis 3, stating that the evolution of niche differences is higher in Páramo lineages, resulting in higher plant species richness in the Páramo than in the Afroalpine, could neither be confirmed nor rejected. Previous research has shown that species in tropical alpine ecosystems exhibit a high degree of phylogenetic clustering^53^, and that pre-adaptation to temperate climates seems necessary for species establishment^20,21^. We found that once a lineage has colonised tropical alpine ecosystems, species appear to successfully adapt and speciate in the available niche space, as shown by the low degree of niche overlap between species of the same lineages (Supplementary Fig. E9; Supplementary Table S5). The degree of newly occupied niche space increases with genetic distance for four of the six studied lineages (Fig. 4A). For three lineages, namely *Loricaria*, *Senecio* (both Páramo), and *Dendrosenecio* (Afroalpine), niche differentiation is correlated with distance between ranges, indicating niche divergence due to geographic separation and different associated climatic conditions. For *Lobelia, Helichrysum* (both Afroalpine) and *Oritrophium* (Páramo), genetic distance does not correlate with range distance nor with new niche space, indicating that niche divergence more likely results from ecological differentiation. Hence, the higher species richness in the Páramo cannot be solely attributed to the evolution of new climatic niche conditions.

Taken together, the results suggest that niche divergence in the Páramo is driven by range overlap (result of hypothesis 2) rather than purely genetic or geographic distance (result of hypothesis 3). This contrasts with the situation in Andean birds, which exhibit high species richness because of high levels of niche packing^15,54,55^, i.e. high levels of niche overlap in range-overlapping species. An increase in species richness via niche packing is often associated with divergence in a non-climatic niche dimension, for example, diet and foraging in tropical birds^54^, or shifts in pollinators and flowering times^56^. In the Afroalpine, the higher niche overlap of range-overlapping species suggests that competition might be higher, leading to lower species richness due to higher chances of species extinction or local exclusion. These findings indicate that niche partitioning to reduce competition might play a more significant role in shaping biodiversity patterns in the Páramo than in the Afroalpine.

The emerging pattern for tropical alpine ecosystems suggests that the differences in species richness between the South American Andes and the East African mountains are due to differences in the persistence and differential *in situ* adaptation of lineages in these diverse and ever-changing environments. The species richness differences between the tropical alpine Andes (higher) and tropical alpine Africa (lower) can likely be attributed to different means of resource partitioning: while niche separation prevails in the Páramo for range-overlapping species, niche packing is prevalent in the Afroalpine. In the northern Andes, with their vast, nearly contiguous mountain chains, climatic oscillations during the Pleistocene resulted in cycles of species and range fragmentation and merging^24^, thereby likely promoting pulses of secondary contact of populations^3^, which might have promoted niche separation. Coupled with a lower extinction threat than in the Afroalpine due to higher habitat stability from the Last Glacial Maximum to the present^17^, this has likely contributed to pronounced niche separation and, consequently, a more species-rich Páramo flora^57^. In contrast, the Afroalpine populations perched on isolated mountain tops, each with a relatively small area and lacking habitat connectivity during the Pleistocene^17,23,58^.

Overall, the complex niche and range patterns observed for the two tropical alpine ecosystems in our study underscore the need for nuanced and region-specific approaches to understanding and conserving biodiversity. Our study also highlights the importance of considering lineage-specific responses to ecological pressures. The distinct patterns across lineages indicate that multiple evolutionary forces are simultaneously at play, with varying intensities depending on the time, region, and community, together shaping species richness patterns. This study represents a major step forward in unravelling the complex evolutionary forces and ecological pressures shaping these remarkable ecosystems and underscores the importance of further investigations to understand the formation of biodiversity hotspots.

## Material and Methods

### Phylogenetic relationships and genetic distances

We selected six characteristic flowering plant lineages that are common in the tropical alpine ecosystems: *Oritrophium*, *Senecio*, and *Loricaria* as representatives of the Páramo, and *Lobelia, Helichrysum,* and *Dendrosenecio* as representatives of the Afroalpine. Except for *Lobelia*, we relied on recently published species trees reconstructed using target enrichment (Hyb-Seq) data^29–31,34,35^. For *Oritrophium* and a tropical alpine clade of *Helichrysum* published phylogenies were expanded. For *Senecio*, we recalculated the phylogeny to improve missing data. For the giant lobelias, we calculated a new species tree based on the same Hyb-Seq protocols for data generation and analysis. For sampling details, sequencing, and phylogenetic reconstruction, see Supplementary Note S2 and Supplementary Table S7-S9.

We used phylogenetic estimates with single representatives of species that showed no signature of introgression for further analyses. We rendered the six trees ultrametric using a penalised likelihood approach implemented in treePL v.1.0^36,37^, based on secondary calibration points from dated phylogenies^19,31,38,40,59,60^ (Supplementary Table S10). Automatic estimation of the optimal parameters for cross-validation was done using a wrapper script (https://github.com/tongjial/treepl_wrapper). The resulting ultrametric trees were analysed for pairwise genetic distance among species in each lineage using the function cophenetic.phylo in ape v.5.7 in R v.4.2.1^61,62^.

### Occurrence records

Occurrence records were mainly obtained from museum specimens’ label information to obtain a set of highly curated data with confirmed taxonomic identity (for details see Supplementary Data S1) and supplemented with GBIF data (querying species names) for range-restricted and rare species. Occurrences were cleaned by filtering duplicate records (identical coordinates) per species, and filtering records outside temperate climates within the tropics as defined in ^17^, as all species are reported to be exclusively montane and/or alpine. Species represented by ≤4 records were omitted. We used all occurrences obtained after filtering (5 to 295 occurrences per species) in the subsequent analyses to make use of the full data for narrowly endemic species.

### Geographic range and climatic niche approximation

Range and niche metrics were calculated in R using functions of the packages sf v.1.0-12, raster v.3.6-14, concaveman v.1.1.0, ecospat v.3.3, ade4 v.1.7-22, and rgeos v.0.6-1^63–68^. To approximate species’ ranges in geographic space, we first calculated spatial polygons around occurrence records of each species (the extent of occurrence), drawn as convex hull around occurrences with a buffer of one using the concaveman function. Within these polygons, we filtered regions outside temperate climates^17^ to better approximate the area of occupancy of the species. Removing non-temperate climatic regions is especially important in Africa, where the high altitude area is divided by extensive lowlands which would significantly inflate range sizes. For species range size, we summarised the spatial extent of species ranges in km^2^. Pairwise range overlap among species per lineage was approximated by the percentage of range overlap of the smaller-ranged species^69^. For non-overlapping species, pairwise range distance was approximated by the minimum distance between species ranges (in m; zero in range-overlapping species).

To approximate species’ climatic niche, we relied on present climate data from CHELSA v.1.2 (30s resolution)^41^ following the method by Broenniman et al. ^42^. We first constructed a global tropical alpine environment using principal component analysis (PCAenv) based on 62,184 spatial points randomly sampled from the area of occupancy of all studied species and the selected eight bioclimatic variables (Supplementary Note S3). For niche size, we multiplied the variances of values of principal components (individuals’ scores) of the first two axes^70^. For niche overlap, we calculated the pairwise similarity metric Schoener’s *D*^55^ corrected by the ratio of the kernel density distribution of available climates among regions^42^. We further quantified niche dynamics among species by calculating metrics for niche stability, niche unfilling, and niche expansion^71^, summarising the two latter as ‘new niche space’ (sum of expansion plus unfilling and then divided by two).

### Statistical analysis

To test our first hypothesis, we tested if niche size and range size between regions and lineages were significantly different using the Wilcoxon rank-sum two-sample test (nonparametric t-test) and Kruskal-Wallis test, respectively. To test for a positive correlation of niche size and range size, as shown before^45,46^, we fitted a PGLS model of niche size∼range size for each lineage based on the R package caper v.1.0.1^72^. To test if niche size correlates with the variability of PC1 or PC2, we fitted another PGLS of niche size∼niche size-PC1 and niche size∼niche size-PC2. For the second hypothesis, we tested if niche overlap as well as range overlap between regions and lineages were significantly different using the same statistical tests as above. Due to spatial autocorrelation, niche and range overlap are likely correlated, which was tested using a linear regression model y = a + bx, where y is the response variable, here niche overlap, and x is the predictor variable, here range overlap, with the slope b, and intercept a. The third hypothesis was tested using four linear regression models y = a + bx. The predictor variable, x, is here genetic distance in million years. As response variable y, we used range distance, the degree of new niche space, and distances between PC axis optima.

Scripts developed for this manuscript are publicly available under: https://github.com/mkandziora/NicheSizeDifferences.

## Data sharing plans

Code to perform the niche and range analyses can be found under https://github.com/mkandziora/NicheSizeDifferences. Sequence data used in this study can be found in Genbank SRA under Bioproject number: PRJNA1092049 (https://www.ncbi.nlm.nih.gov/bioproject/?term=PRJNA1092049). Alignments, phylogenetic trees and used occurrence data are available as Supplementary.

## Acknowledgment

We thank Sileshi Nemomissa and Sebsebe Demissew (Ethiopia), Geoffrey Mwachala (Kenya), Pantaleo Munishi (Tanzania), and Gerald Eilu (Uganda) for assistance and for facilitating collection permits for the AFROALP-II and SpeciationClock projects from the Ethiopian Wildlife Conservation Authority, the National Museums of Kenya, the Tanzania Commission for Science and Technology, the Tanzania National Parks Authority, the Uganda National Council for Science and Technology (UNCST), and the Uganda Wildlife Authority (UWA). Collection in Rwanda was done under Research permit No NCST/482/304/2022 and with the permission of Rwanda Development Board (RDB) Tourism and Conservation to collect in the Volcanoes National Park. We thank Prof. Elias Bizuru for allowing affiliation of MK and MGC to University of Rwanda, Dr. Richard Muvunyi from RDB for his support, the staff of the Volcanoes National Park for field assistance; and Beth Kaplin and the staff from the National Herbarium of Rwanda, as well as the staff of the Ellen DeGeneres Campus in Musanze, for logistic support. Collections in Ecuador were supported by a research permit issued by the Ministerio del Ambiente, Quito, Ecuador (No. 09-IC-FLO- DNB/MAE, 24-2010-IC-FLO-DPAP-MA, MAE-DNB-CM-2018-0082). Computational resources were supplied by the project “e-Infrastruktura CZ” (e-INFRA LM2018140) provided within the program Projects of Large Research, Development and Innovations Infrastructures. We also thank Juan Manuel Gorospe for providing the phylogenetic trees of *Dendrosenecio* and *Oritrophium*.

## Authors contributions

Conceptualization: MK, NN; Methodology: MK; Formal analysis: MK; Investigation: MK, DLAV; Resources: PS, CB, AG, MK, MGC, LG, DC; Data Curation: MK; Writing - Original Draft: MK, RS, NN, CB; Writing - Review & Editing: MK, DLAV, CB, AG, MGC, FK, PS, NN, RS; Visualisation: MK; Supervision: RS, NN; Project administration: MK, RS; Funding acquisition: MK, RS, FK, MGC, CB

