## Supplementary Material for "Different resource partitioning explains plant species richness patterns in tropical alpine ecosystems"

This Supplementary Material consists of:

Supplementary Notes S1 to S3
Figures S1 to S13
Tables S1 to S10
Supplementary Data S1

*Supplementary Note S1. Phylogenetic reconstruction of* Oritrophium, Senecio, Helichrysum, *and giant lobelias*

We opted to improve the phylogenetic sampling of the tropical alpine genera of *Oritrophium*, *Senecio*, *Helichrysum*, and giant lobelias. Previous research highlighted the non-monophyly of *Oritrophium*, with some species showing closer affinities to *Erigeron*[1]. However, our analysis confirms that *Oritrophium* s.str. is well supported as a monophyletic clade (local posterior probability [LPP] = 1 and bootstrap support [BS] = 98%; Fig. S1). Compared to the existing Hyb-Seq phylogeny of *Oritrophium*[2], in our extended version, *O. cocuyense* (Cuatrec.) Cuatrec. was excluded here due to introgression. Further, our phylogenies, based on an ASTRAL v.5.7.7[3] species tree as well as a concatenated RAxML-NG v.0.9.0[4] tree, reveal the Mexican *O. orizabense* G.L.Nesom as sister to the remaining species, contradicting an earlier hypothesis that is based on Sanger sequencing data[1]. In that previous study[1], *O. hieracioides* Wedd. was the earliest branching species, followed by *O. orizabense* and *O. ferrugineum* Wedd.; *O. hieracioides* and *O. ferrugineum* form the next basal clade in our analyses (Fig. S1). The widespread species *O. limnophilum* (Sch. Bip.) Cuatrec. is sister to a clade comprising the remaining species. Those remaining species are found in two main clades, one comprising *O. aciculifolium* Cuatrec., *O. yacuriense* Arnelas & J.Calvo, and *O. repens* (Kunth) Cuatrec., and the other clade being composed of *O. ollgardii* Cuatrec., *O. mucidum* Cuatrec., *O. llanganatense* Sklenár & H.Robinson, *O. peruvianum* (Lam.) Cuatrec., and *O. crocifolium* (Kunth) Cuatrec.

For our *Senecio* phylogeny, built upon previous work[5], we replaced some samples to reduce missing data in the alignments; sampling information is provided in Supplementary Table S7-S8. The resulting phylogeny was similar in resolution and support to the published phylogeny (Fig. S2); the series itself is well supported (LPP = 1/BS = 100%). Compared to the previous Hyb-Seq phylogeny[5], we excluded six samples for the single-tip phylogeny here due to signals of introgression of greater than 10% (Fig. S7; *S. patens* (Kunth) DC., *S. puracensis* Cuatrec., *S. superparamensis* Sklenář, *S. expansus* Wedd., *S. longepenicillatus* Sch.Bip. ex Wedd and *S. lingulatus* (Schltdl.) Cuatrec.). Overall support and resolution remained similar. Especially internal nodes are not supported, which resulted in a different resolution at the base.

For the *Helichrysum* phylogeny, our analysis confirms the well-supported afroalpine clade (1 LPP = 1, BS = 100%; Fig. S3; part of TA1 in [6], with no major differences in resolution and support). The clade is composed of *H. stuhlmannii* O. Hoffm., *H. amblyphyllum* Mattf., *H. ellipticifolium* Moeser, *H. formosissimum* Sch.Bip., and *H. meyeri-johannis* Engl. in clade 1, and *H. argyranthum* O.Hoffm., *H. nandense* S.Moore, *H. brownei* S.Moore, *H. newii* Oliv. & Hiern, and *H. chionoides* Philipson in clade 2. Most species form monophyletic clades, with the exception of *H. meyeri-johannis*, which is nested within *H. formosissimum* (without support), and of *H. nandense,* that is nested within *H. argyranthum.* Some authors consider *H. nandense* as a synonym of *H. argyranthum*, which would explain these results.

For the newly calculated giant lobelia phylogeny, we used the herbal species of *Lobelia lindblomii* Mildbr. as outgroup taxon. We also included the giant West African *L. columnaris* Hook f. in the analysis as well as *L. thuliniana* E.B.Knox, a species occurring south of the tropical alpine regions. The clade of giant lobelias is well supported (Fig. S4; 1 LPP, 100% BS) and reveals the monophyly of most species. *Lobelia columnaris* is the most basal species, followed by a clade of the widespread montane *L. giberroa* and *L. thuliniana* (clade 3). One sample of *L. giberroa* shows a different placement in the concatenated phylogeny compared to the ASTRAL species tree (sample highlighted with a pink star in Fig. S4; within the ASTRAL species tree it falls together with the other *L. giberroa* samples). Sister to clade 3 are two subclades: clade 1 comprises the two Ethiopian species of *L. rhynchopetalum* Hemsl. and *L. acrochila* (E.Wimm.) E.B.Knox, and the species from the eastern branch, *L. aberdarica* R.E. & T.C.E.Fries, *L. deckenii*, *L. bambuseti* R.E.Fr. & T.C.E.Fr., and *L. telekii* Schweinf. Clade 2 comprises the species from the western branch of the East African Rift, *L. stuhlmannii* Schweinf. ex Stuhlmann, *L. wollastonii* Baker f., and *L. deckenii* subsp. *bequaertii* De Wild.

We included at least two samples per species to confirm monophyly of the species, with the exception of *L. thuliniana and L. columnaris* where only one sample was available. The widespread species *L. deckenii* (Asch.) Hemsl. was confirmed to be monophyletic (subsp. *gregoriana*, subsp. *burtii*, subsp. *deckenii*), with the exception of *L. deckenii* subsp. *bequaertii*, which is found in a clade together with other species that occur along the western branch of the African Rift. The species *L. mildbraedii* Engl. is not monophyletic; two samples were found in clade 1 and clade 2 each, while all samples have been collected in south-western Uganda, similar to the main distribution of species in clade 2. For the single-tip phylogeny (Fig. S10), we excluded duplicated samples, with the exception of the non-monophyletic *L. mildbraedii* which is represented by two samples.

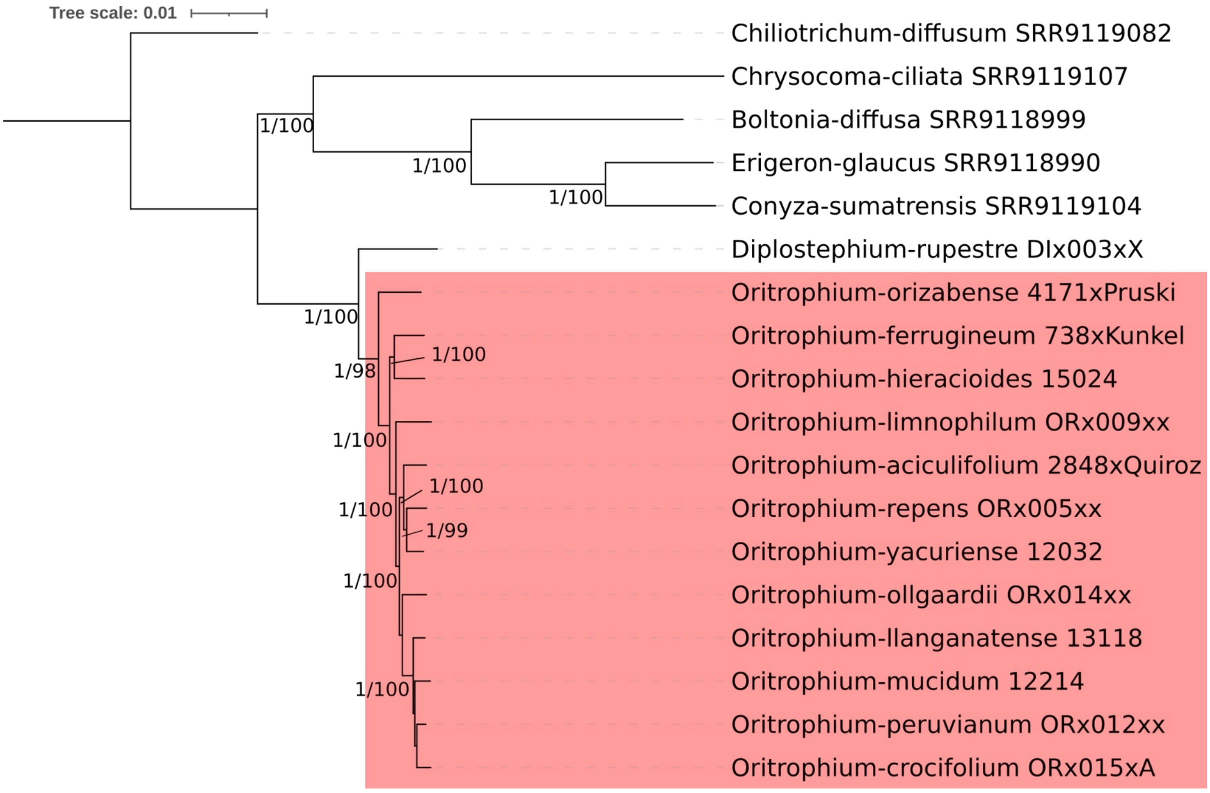

Figure S1: Best Maximum Likelihood (ML) phylogeny of the South American tropical alpine genus *Oritrophium* s.str. based on Hyb-Seq exons. The phylogeny was reconstructed using RAxML-NG. Values at branches show support based on ASTRAL local posterior probability/ML bootstrap greater 0.94/94.

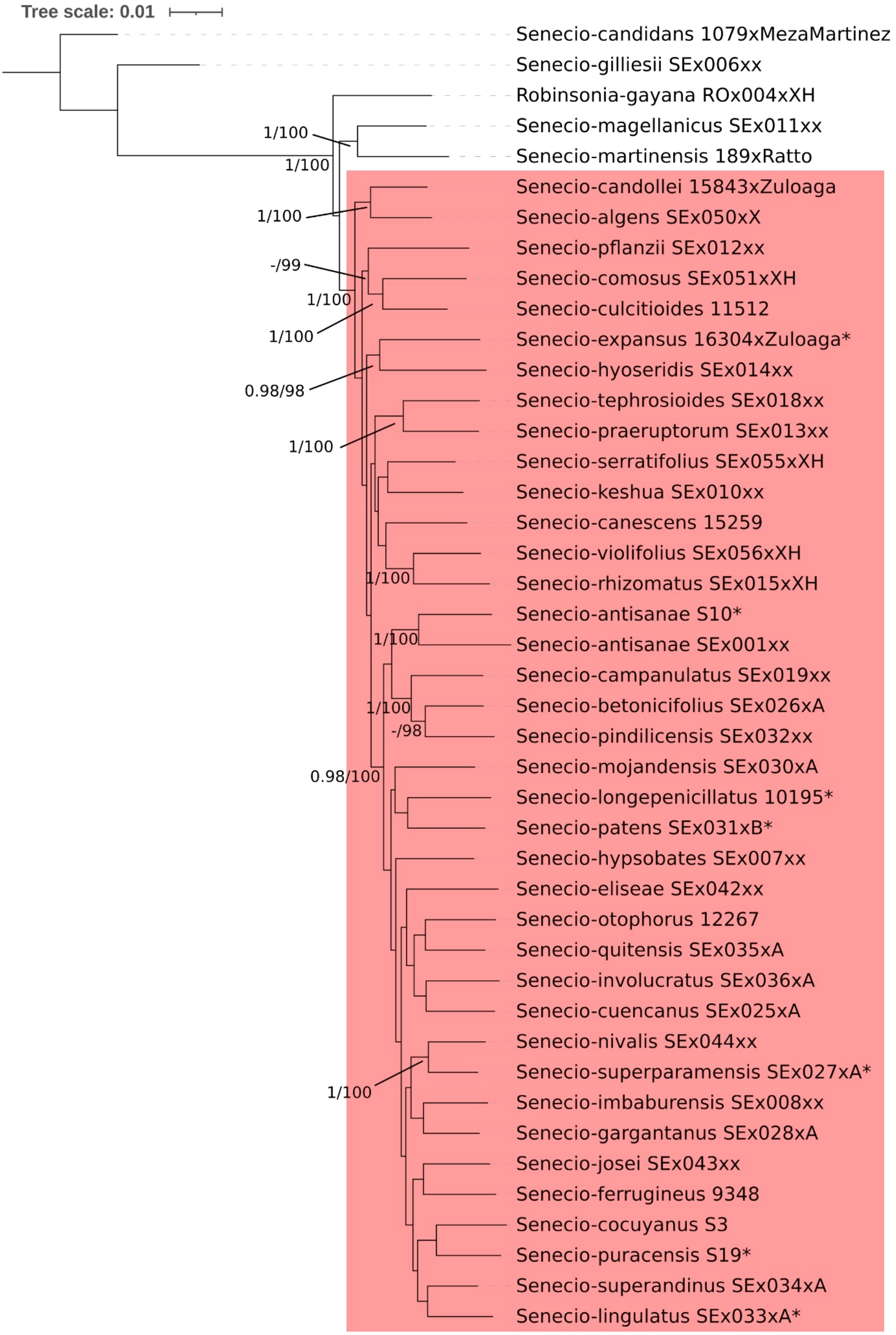

Figure S2. Best Maximum Likelihood (ML) phylogeny of the South American clade of *Senecio* ser. *Culcitium* based on Hyb-Seq exons. The phylogeny was reconstructed using RAxML-NG. Values at branches show support based on ASTRAL local posterior probability/ML bootstrap greater 0.94/94. Asterisks show samples that were removed for the single-tip ultrametric phylogeny.

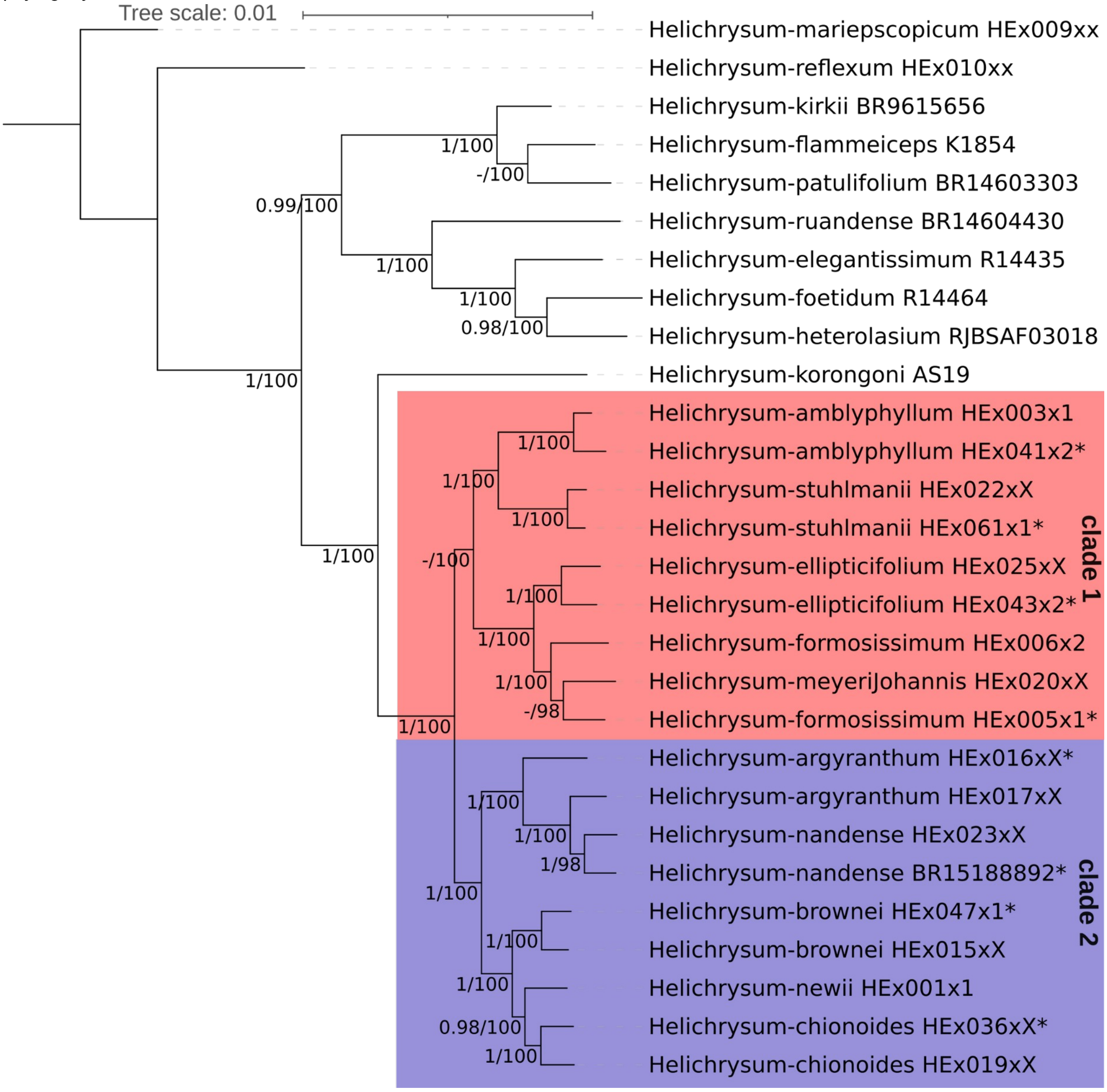

Figure S3: Best Maximum Likelihood (ML) phylogenies of the East African tropical alpine *Helichrysum* clade based on Hyb-Seq exons. The phylogeny was reconstructed using RAxML-NG. Values at branches show support based on ASTRAL local posterior probability/ML bootstrap greater 0.94/94. Asterisks show samples that were removed for the single-tip ultrametric phylogeny.

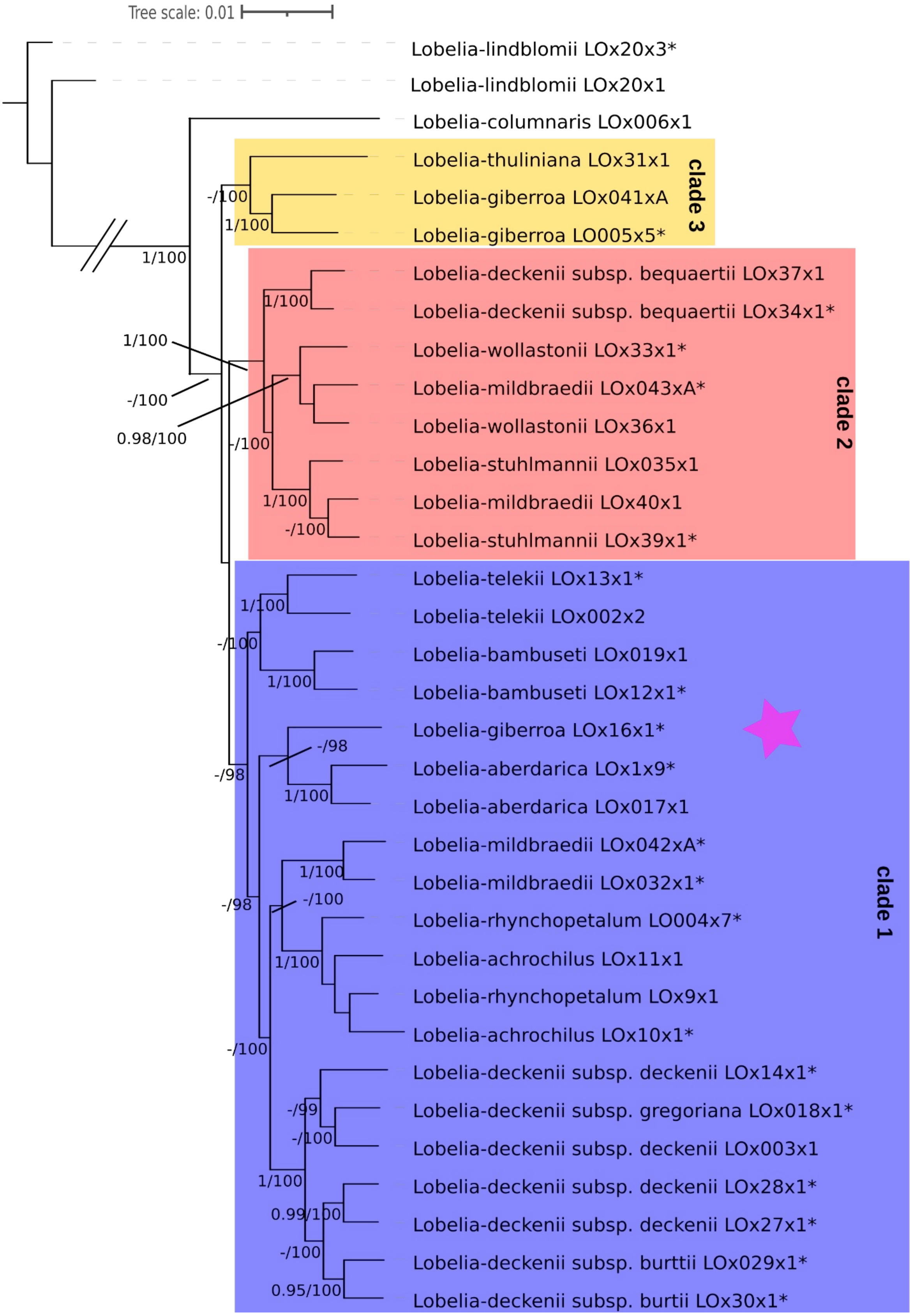

Figure S4: Best Maximum Likelihood (ML) phylogeny of the East African giant *Lobelia* clade based on Hyb-Seq exons. The phylogeny was reconstructed using RAxML-NG. Values at branches show support based on ASTRAL local posterior probability/ML bootstrap greater 0.94/94. Single asterisks indicate samples that were removed for the single-tip ultrametric phylogenies, pink star indicates samples that are placed differently in the ASTRAL species tree. Double diagonal line indicates that the branch was shortened for display.

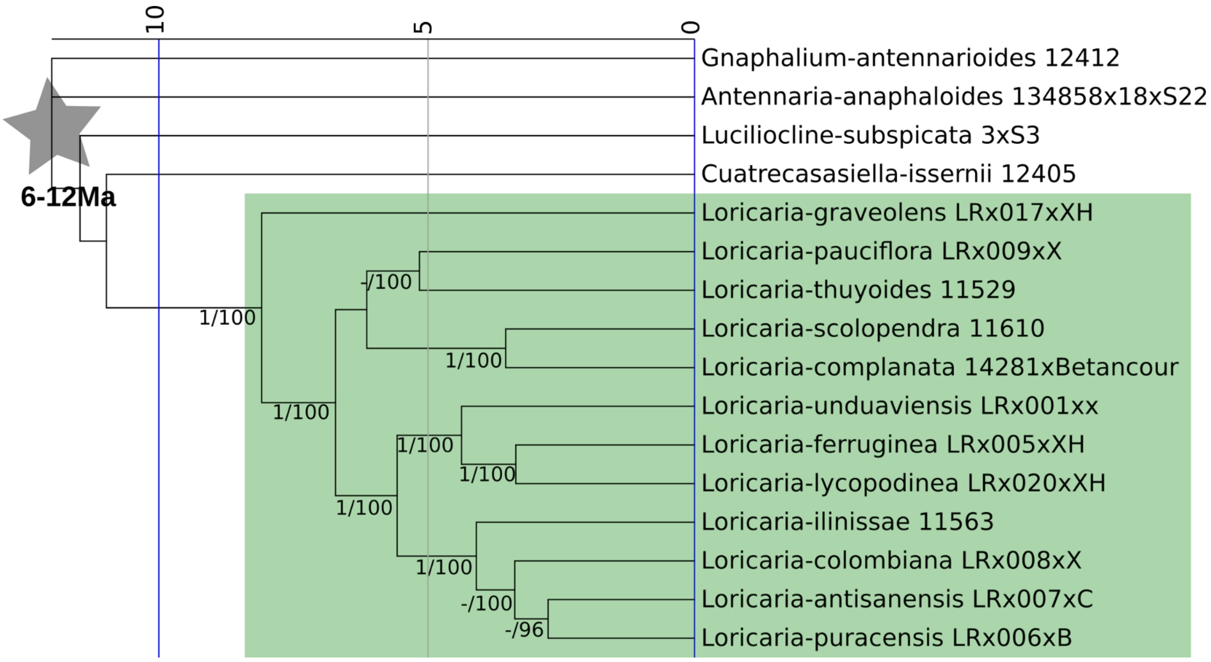

Figure S5. Ultrametric phylogeny of the South American tropical alpine genus *Loricaria* including outgroup samples, based on the best Maximum Likelihood (ML) RAxML-NG phylogeny and a penalised likelihood approach. Values at branches show support based on ASTRAL local posterior probability/ML bootstrap greater 0.94/94. Stars point to nodes used as secondary calibration points with the respective constraints. Timescale in million years (Ma) shown at the top. Green box indicates tropical alpine clade.

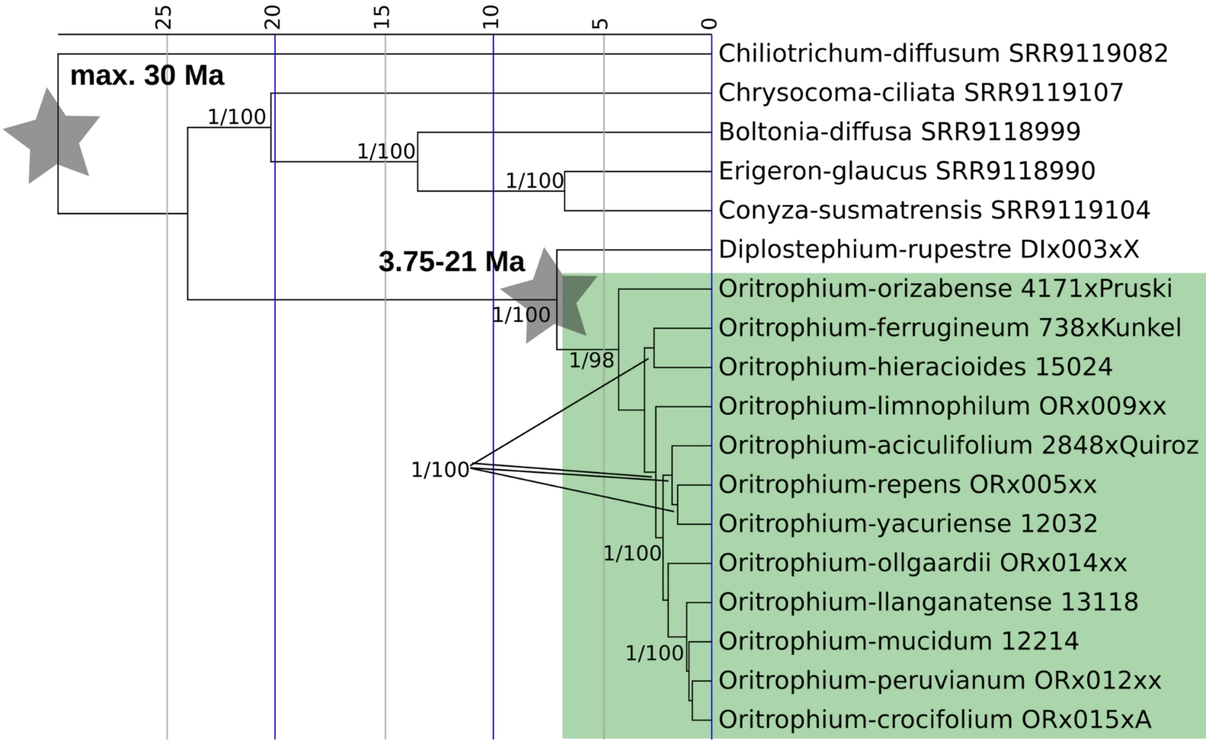

Figure S6. Ultrametric phylogeny of the South American tropical alpine genus *Oritrophium* including outgroup samples, based on the best Maximum Likelihood (ML) RAxML-NG phylogeny and a penalised likelihood approach. Values at branches show support based on ASTRAL local posterior probability/ML bootstrap greater 0.94/94. Stars point to nodes used as secondary calibration points with the respective constraints. Timescale in million years (Ma) shown at the top. Green box indicates tropical alpine clade.

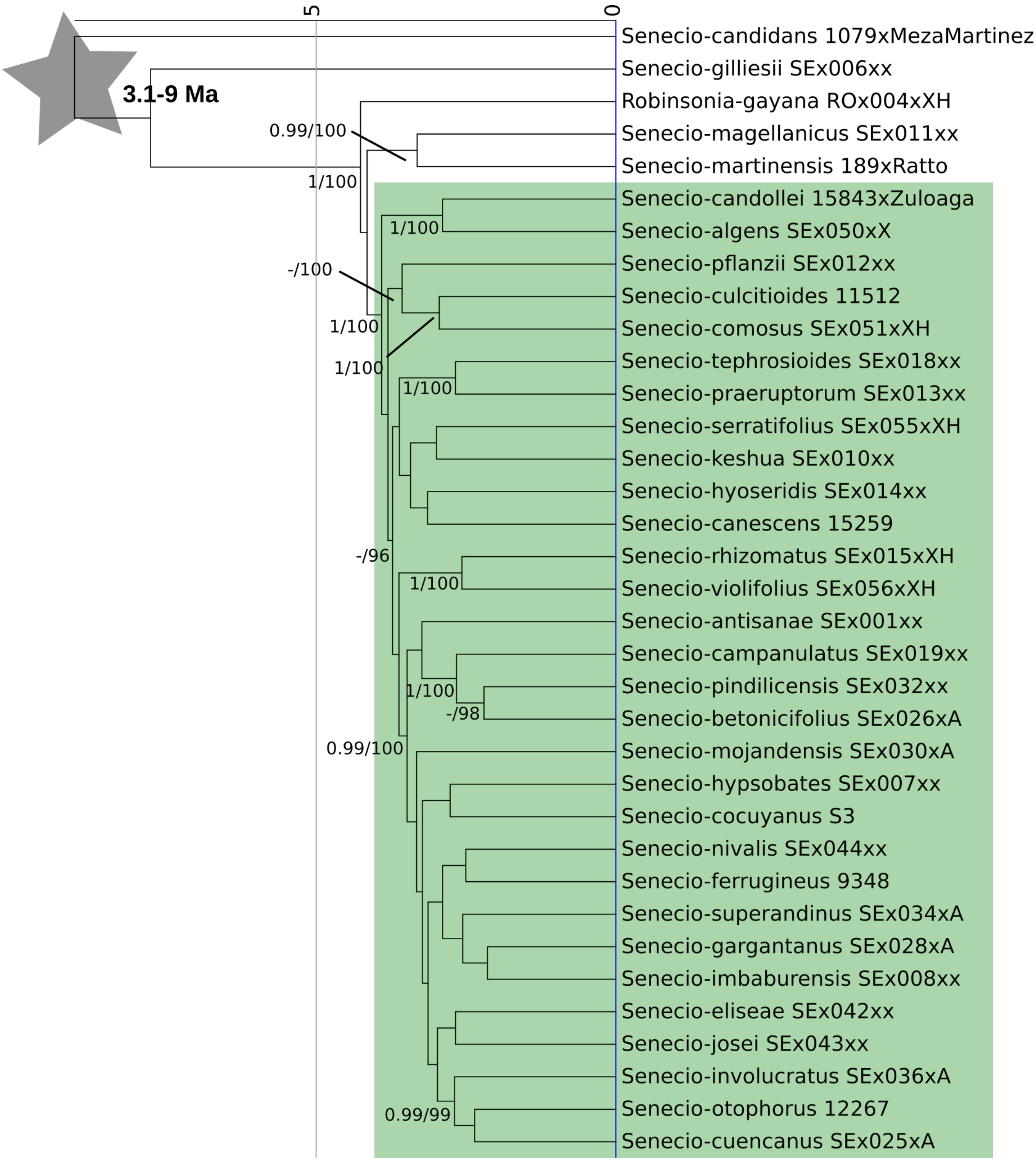

Figure S7. Ultrametric phylogeny of the South American tropical alpine lineage *Senecio* ser. Culcitium including outgroup samples, based on the best Maximum Likelihood (ML) RAxML-NG phylogeny and a penalised likelihood approach. Values at branches show support based on ASTRAL local posterior probability/ML bootstrap greater 0.94/94. Stars point to nodes used as secondary calibration points with the respective constraints. Timescale in million years (Ma) shown at the top. Green box indicates tropical alpine clade.

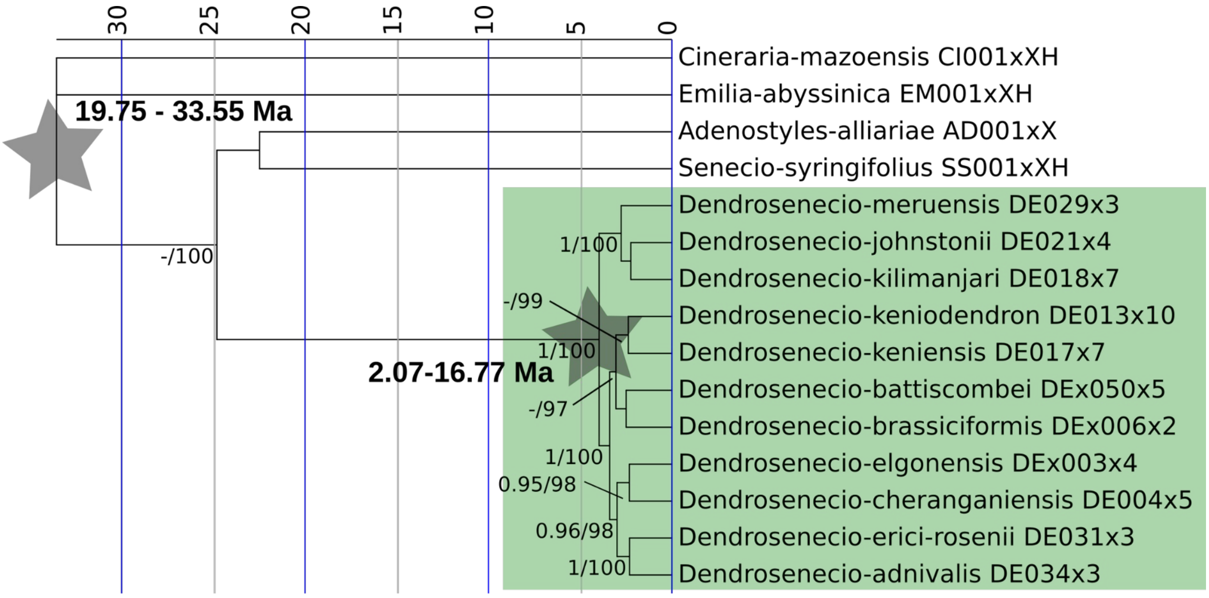

Figure S8. Ultrametric phylogeny of the East African tropical alpine genus *Dendrosenecio* including outgroup samples, based on the best Maximum Likelihood (ML) RAxML-NG phylogeny and a penalised likelihood approach. Values at branches show support based on ASTRAL local posterior probability/ML bootstrap greater 0.94/94. Stars point to nodes used as secondary calibration points with the respective constraints. Timescale in million years (Ma) shown at the top. Green box indicates tropical alpine clade.

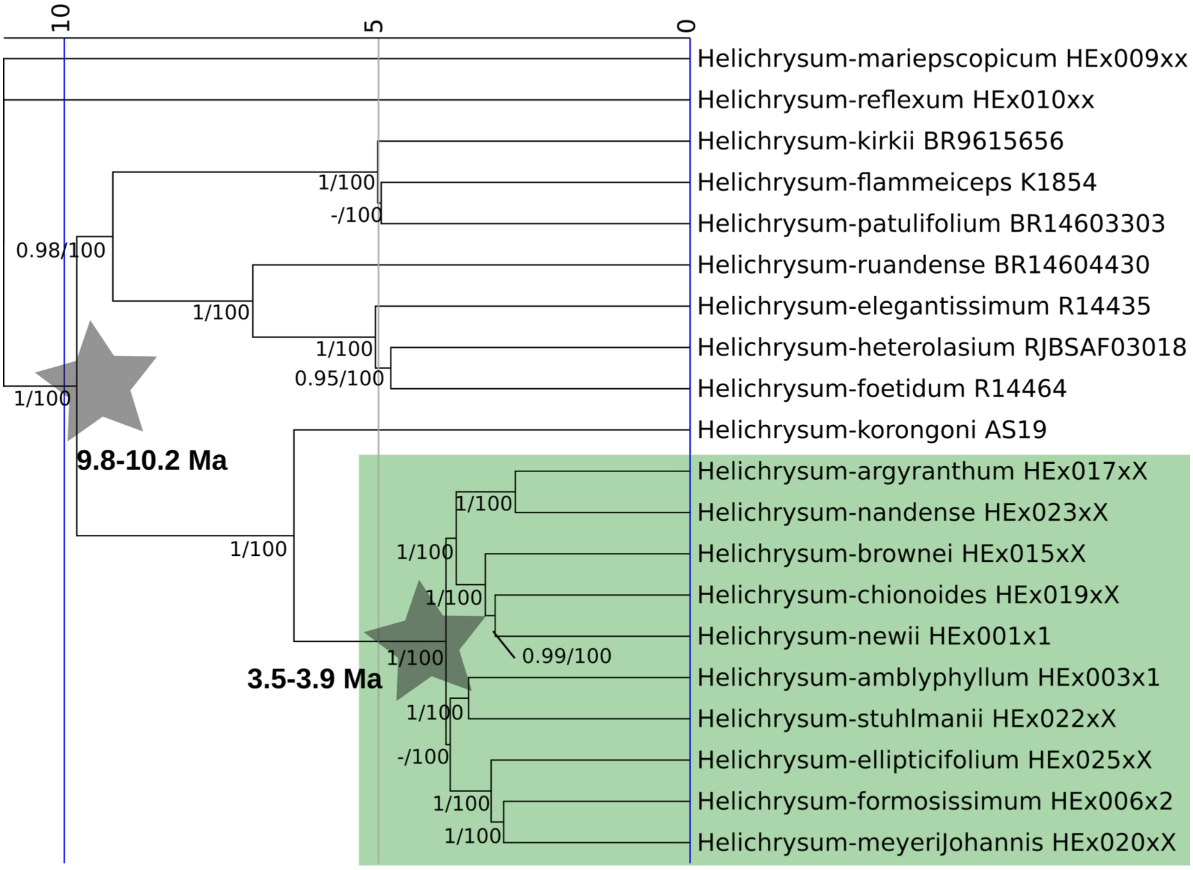

Figure S9. Ultrametric phylogeny of the East African tropical alpine clade of *Helichrysum*, including outgroup samples, based on the best Maximum Likelihood (ML) RAxML-NG phylogeny and a penalised likelihood approach. Values at branches show support based on ASTRAL local posterior probability/ML bootstrap greater 0.94/94. Stars point to nodes used as secondary calibration points with the respective constraints. Timescale in million years (Ma) shown at the top. Green box indicates tropical alpine clade.

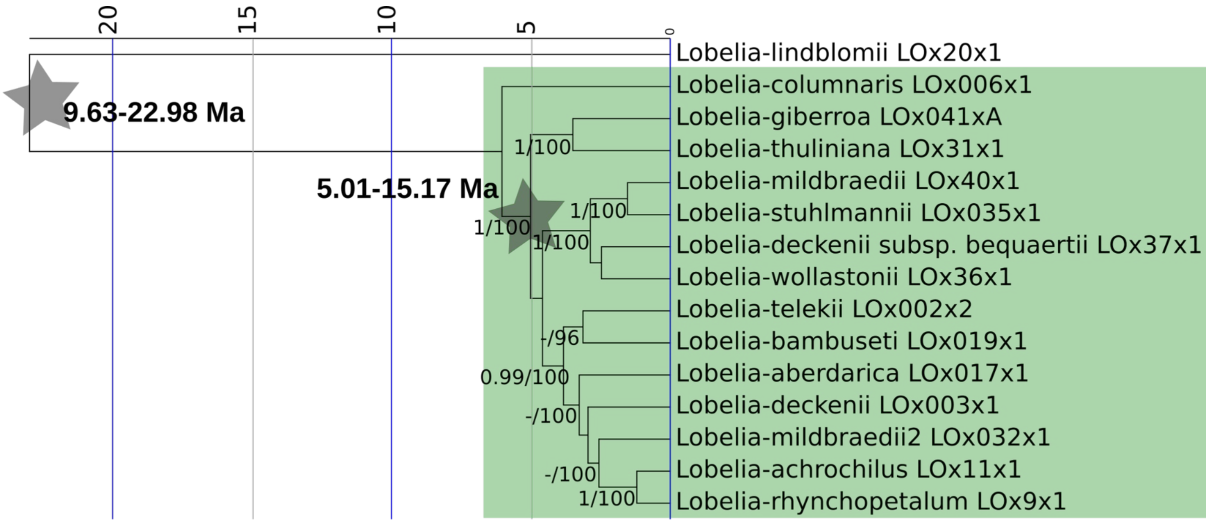

Figure S10. Ultrametric phylogeny of the African tropical alpine giant *Lobelia* clade including outgroup samples, based on the best Maximum Likelihood (ML) RAxML-NG phylogeny and a penalised likelihood approach. Values at branches show support based on ASTRAL local posterior probability/ML bootstrap greater 0.94/94. Stars point to nodes used as secondary calibration points with the respective constraints. Timescale in million years (Ma) shown at the top. Green box indicates tropical alpine clade.

Table S1. Molecular age estimates in millions of years and number of accepted species for the different lineages.

| | number of species | crown age | stem age |
|---|---|---|---|
| *Dendrosenecio* | 11 | 3.96 | 25.31 |
| *Lobelia* | 13 | 6.03 | 22.98 |
| *Helichrysum* | 10 | 3.9 | 6.33 |
| *Loricaria* | 19 | 7.11 | 10.75 |
| *Oritrophium* | 18 | 4.54 | 7.26 |
| *Senecio* | 36 | 3.89 | 4.14 |

Table S2. Summary of cleaned occurrence data. *highlights species that were supplemented with occurrence points from GBIF. Abbreviations: # - number of; occ - occurrence; spp. - species.

| Eastern African mountain lineages* | | | | | South American Andean lineages | | | | |
|---|---|---|---|---|---|---|---|---|---|
| lineage | species | # occ | mean occ/spp | # spp with occ | lineage | species | # occ | mean occ/spp | # spp with occ |
| *Lobelia* | *aberdarica* | 15 | 37.82 | 11 | *Oritrophium* | *aciculifolium** | 6 | 49.36 | 11 |
| | *acrochila* | 6 | | | | *crocifolium* | 24 | | |
| | *bambuseti* | 14 | | | | *ferrugineum** | 6 | | |
| | *bequaertii* | 15 | | | | *hieracioides* | 15 | | |
| | *deckenii* | 65 | | | | *limnophilum* | 126 | | |
| | *giberroa* | 96 | | | | *marahuacense** | 12 | | |
| | *mildbraedii* | 29 | | | | *ollgaardii** | 10 | | |
| | *rhynchopetalum* | 73 | | | | *orizabense** | 5 | | |
| | *stuhlmannii* | 38 | | | | *peruvianum* | 295 | | |
| | *telekii* | 23 | | | | *repens* | 39 | | |
| | *wollastonii* | 42 | | | | *yacuriense* | 5 | | |
| *Helichrysum* | *argyranthum* | 22 | 31.6 | 10 | *Loricaria* | *antisanensis** | 21 | 36.25 | 12 |
| | *brownei* | 29 | | | | *colombiana* | 18 | | |
| | *chionoides* | 15 | | | | *complanata* | 45 | | |
| | *ellipticifolium* | 11 | | | | *ferruginea* | 27 | | |

| | | | | |
|---|---|---|---|---|
| | *formosissimum* | 95 | | |
| | *guilelmi* | 23 | | |
| | *meyeri-johannis* | 19 | | |
| | *nandense* | 13 | | |
| | *newii* | 53 | | |
| | *stuhlmanii* | 36 | | |
| *Dendrosenecio* | *adnivalis* | 19 | 26.64 | 11 |
| | *battiscombei* | 29 | | |
| | *brassiciformis* | 13 | | |
| | *cheranganiensis* | 18 | | |
| | *elgonensis* | 31 | | |
| | *erici-rosenii* | 36 | | |
| | *johnstonii* | 15 | | |
| | *keniensis* | 30 | | |
| | *keniodendron* | 39 | | |
| | *kilimanjari* | 55 | | |
| | *meruensis* | 8 | | |
| Total | | 1025 | 32.35 | 32 |
| | *graveolens* | 7 | | |
| | *ilinissae* | 68 | | |
| | *leptothamna** | 18 | | |
| | *ollgaardii* | 15 | | |
| | *pauciflora* | 5 | | |
| | *scolopendra* | 20 | | |
| | *thuyoides* | 184 | | |
| | *unduaviensis** | 7 | | |
| *Senecio* | *algens** | 30 | 32.52 | 29 |
| | *antisanae** | 13 | | |
| | *betonicifolius* | 5 | | |
| | *burkartii* | 8 | | |
| | *campanulatus* | 20 | | |
| | *candollei* | 8 | | |
| | *canescens* | 34 | | |
| | *cocuyanus** | 29 | | |
| | *comosus* | 12 | | |
| | *cuencanus* | 7 | | |
| | *culcitioides* | 26 | | |
| | *gargantanus* | 11 | | |
| | *hypsobates* | 7 | | |
| | *imbaburensis* | 10 | | |
| | *involucratus* | 99 | | |
| | *lingulatus* | 56 | | |
| | *longepenicillatus* | 46 | | |
| | *mojandensis* | 28 | | |
| | *nivalis* | 15 | | |
| | *otophorus* | 61 | | |
| | *patens* | 79 | | |
| | *pindilicensis* | 13 | | |
| | *praeruptorum* | 15 | | |
| | *puracensis* | 12 | | |
| | *rhizomatus* | 25 | | |
| | *serratifolius** | 37 | | |
| | *superandinus* | 137 | | |
| | *superparamensis* | 15 | | |
| | *tephrosioides* | 81 | | |
| Total | | 1917 | 39.49 | 52 |

Table S3. Bioclimatic variables used for the calculation of the climatic niche space.

| Bioclim name | Bioclim explanation | Variance inflation factor for log(bioclim) |
|---|---|---|
| bio2 | Mean diurnal air temperature range | 6.94 |
| bio3 | Isothermality, defined as mean diurnal range/temperature annual range | 2.12 |
| bio5 | Maximum temperature of warmest month | 5.25 |
| bio9 | Mean temperature of the driest quarter | 7.48 |
| bio13 | Mean precipitation of the wettest month | 5.17 |
| bio15 | Precipitation seasonality (coefficient of variation) | 6.87 |
| bio18 | Mean precipitation of the warmest quarter | 5.01 |
| bio19 | Mean precipitation of the coldest quarter | 8.27 |

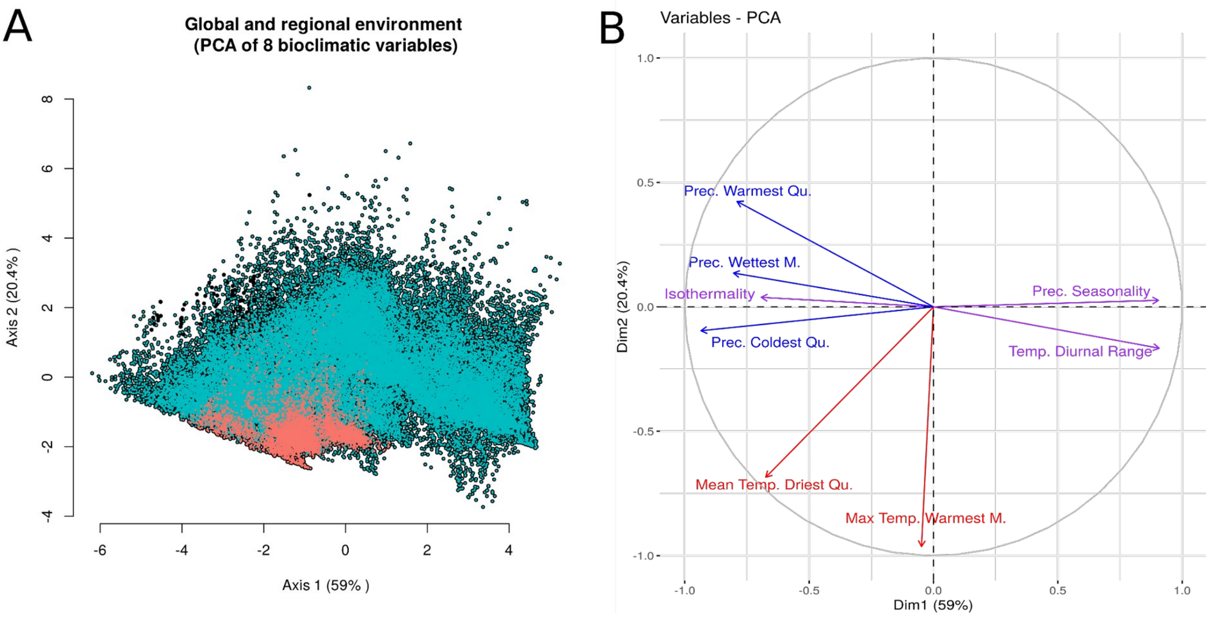

Figure S11: Principal component analysis of the climatic niche space (A) between the two tropical alpine regions (Páramo in light blue and Afroalpine in red) and the contribution of the different bioclimatic variables (B). Red: temperature-related variables; blue: precipitation-related variables; purple: seasonality-related variables.

Table S4. Median values and standard deviation in parenthesis for niche and geographic metrics of the regions. Asterisk and bold font indicate p-value < 0.05 according to the Wilcoxon rank-sum two-sample test. Abbreviations: n - number of data points per subset, Ma - millions of years, PC - principal component, SD - standard deviation.

| | | Afroalpine | Páramo | p-value |
|---|---|---|---|---|
| **niche size *** | median | **0.0910** | **0.286** | **0.01239** |
| | SD | 0.345 | 0.814 | |
| | n | 32 | 52 | |
| **niche size PC axis 1 *** | median | **0.164** | **0.431** | **0.00345** |
| | SD | 0.458 | 0.579 | |
| | n | 32 | 52 | |
| niche size PC axis 2 | median | 0.485 | 0.610 | 0.1185 |
| | SD | 0.308 | 0.481 | |
| | n | 32 | 52 | |
| PC axis 1 optima | median | 1.63 | 1.63 | 0.723 |
| | SD | 0.521 | 1.15 | |
| | n | 32 | 52 | |
| **PC axis 2 optima *** | median | **5.18** | **3.98** | **2.987e-08** |
| | SD | 0.679 | 0.788 | |
| | n | 32 | 52 | |
| **PC axis 1 distance *** | median | **0.565** | **1.02** | **2.496e-12** |
| | SD | 0.468 | 1.11 | |
| | n | 155 | 527 | |
| PC axis 2 distance | median | 0.722 | 0.782 | 0.1737 |
| | SD | 0.588 | 0.614 | |
| | n | 155 | 527 | |
| **niche overlap [%] of…** | | | | |
| …all taxa | median | 0.084 | 0.082 | 0.7179 |
| | SD | 0.205 | 0.164 | |
| | n | 155 | 527 | |
| **…range overlapping taxa [>0.1%] *** | median | **0.227** | **0.156** | **0.01422** |
| | SD | 0.231 | 0.186 | |
| | n | 73 | 246 | |
| **…**non-overlapping [<=0.1%] | median | 0.0185 | 0.032 | 0.1155 |
| | SD | 0.0914 | 0.117 | |
| | n | 82 | 271 | |
| **range size * [km$^2$]** | median | **2981** | **30361** | **1.51e-05** |
| | **SD** | 21008 | 102536 | |
| | **n** | 32 | 51 | |
| range overlap for all taxa [%] | median | 0.00698 | 0.0218 | 0.9387 |
| | SD | 0.372 | 0.360 | |
| | n | 155 | 517 | |

| | | Afroalpine | Páramo | p-value |
|---|---|---|---|---|
| range overlap for species that overlap [>0km2; %] | median | 0.617 | 0.633 | 0.6704 |
| | SD | 0.280 | 0.265 | |
| | n | 73 | 246 | |
| | median | 525 | 569 | 0.3559 |
| range distance between non-overlapping taxa [km] | SD | 342 | 609 | |
| | n | 76 | 245 | |
| crown age [Ma] | median | 3.96 | 4.54 | 1 |
| | SD | 1.21 | 1.7 | |
| | n | 3 | 3 | |
| | median | **7.80** | **7.42** | **0.0002475** |
| **genetic distance [Ma] *** | SD | 1.43 | 1.88 | |
| | n | 146 | 343 | |

Table S5. Median values and standard deviation for niche and geographic metrics of the different lineages. Asterisk and bold font indicate p-value < 0.05 according to the Kruskal-Wallis test. Abbreviations: n - number of data points per subset, PC - principal component, SD - standard deviation.

| lineage | | *Dendro-senecio* | *Heli-chrysum* | *Lobelia* | *Loricaria* | *Oritrophium* | *Senecio* | p-value |
|---|---|---|---|---|---|---|---|---|
| | Median | **0.0299** | **0.271** | **0.118** | **0.239** | **0.108** | **0.579** | **0.00135** |
| **niche size *** | SD | 0.0423 | 0.414 | 0.370 | 0.476 | 0.628 | 0.935 | |
| | n | 11 | 10 | 11 | 12 | 11 | 29 | |
| niche overlap… | | | | | | | | |
| **…of all taxa** | Median | **0.027** | **0.275** | **0.043** | **0.0255** | **0.015** | **0.1** | **1.146e-13** |
| | SD | 0.151 | 0.219 | 0.186 | 0.136 | 0.126 | 0.170 | |
| | n | 55 | 45 | 55 | 66 | 55 | 406 | |
| | Median | **0.251** | **0.28** | **0.115** | **0.102** | **0.054** | **0.18** | **5.755e-05** |
| **…of range overlapping taxa [>0.1]*** | SD | 0.270 | 0.218 | 0.225 | 0.169 | 0.158 | 0.188 | |
| | n | 6 | 41 | 26 | 29 | 21 | 196 | |
| | Median | **0.021** | **0.092** | **0.004** | **0.003** | **0.0015** | **0.0485** | **0.001419** |
| **…non-overlapping[<=0.1] *** | SD | 0.0828 | 0.0816 | 0.107 | 0.0721 | 0.1061 | 0.122 | |
| | n | 49 | 4 | 29 | 37 | 24 | 210 | |
| | Median | **0.0892** | **0.466** | **0.234** | **0.371** | **0.313** | **0.524** | **0.0002054** |
| **PC axis 1 size*** | SD | 0.0559 | 0.385 | 0.624 | 0.337 | 0.490 | 0.652 | |
| | n | 11 | 10 | 11 | 12 | 11 | 29 | |
| | Median | **0.333** | **0.639** | **0.414** | **0.493** | **0.346** | **0.878** | **0.03026** |
| **PC axis 2 size*** | SD | 0.197 | 0.294 | 0.350 | 0.418 | 0.430 | 0.491 | |
| | n | 11 | 10 | 11 | 12 | 11 | 29 | |
| | Median | -1.58 | -1.63 | -1.69 | -1.69 | -1.30 | -1.80 | 0.9455 |
| PC axis 1 optima | SD | 0.511 | 0.361 | 0.666 | 1.16 | 1.05 | 1.21 | |
| | n | 11 | 10 | 11 | 12 | 11 | 29 | |
| | Median | **5.18** | **5.24** | **5.18** | **4.04** | **4.46** | **3.74** | **1.875e-06** |
| **PC axis 2 optima *** | SD | 0.729 | 0.500 | 0.812 | 0.811 | 0.727 | 0.753 | |
| | n | 11 | 10 | 11 | 12 | 11 | 29 | |
| | Median | **443** | **15835** | **2986** | **22192** | **17353** | **53710** | **2.172e-05** |
| **range size [km2] *** | SD | 1335 | 23903 | 25123 | 38706 | 173759 | 87950 | |
| | n | 11 | 10 | 11 | 12 | 10 | 29 | |
| | Median | **0** | **0.557** | **0.00939** | **0.0000547** | **0.0129** | **0.0357** | **4.462e-10** |
| **range overlap [%] *** | SD | 0.282 | 0.281 | 0.399 | 0.350 | 0.447 | 0.351 | |
| | n | 55 | 45 | 55 | 66 | 45 | 406 | |
| | Median | 0.985 | 0.595 | 0.684 | 0.546 | 0.992 | 0.618 | 0.6699 |
| range overlap of overlapping taxa [>0.1%; %]* | SD | 0.179 | 0.239 | 0.333 | 0.312 | 0.315 | 0.246 | |
| | n | 6 | 41 | 26 | 29 | 21 | 194 | |
| | Median | 484 | 18.4 | 801 | 507 | 563 | 579 | 0.2992 |
| range distance of non-overlapping taxa [km] | SD | 258 | 0.235 | 349 | 709 | 673 | 583 | |
| | n | 47 | 2 | 27 | 33 | 22 | 188 | |
| | Median | **6.77** | **7.80** | **9.16** | **11.5** | **6.16** | **7.42** | **0.000245** |
| **genetic distance [Ma]** | SD | 1.06 | 0.628 | 1.80 | 2.16 | 1.7 | 0.662 | |
| | n | 55 | 36 | 55 | 45 | 45 | 253 | |

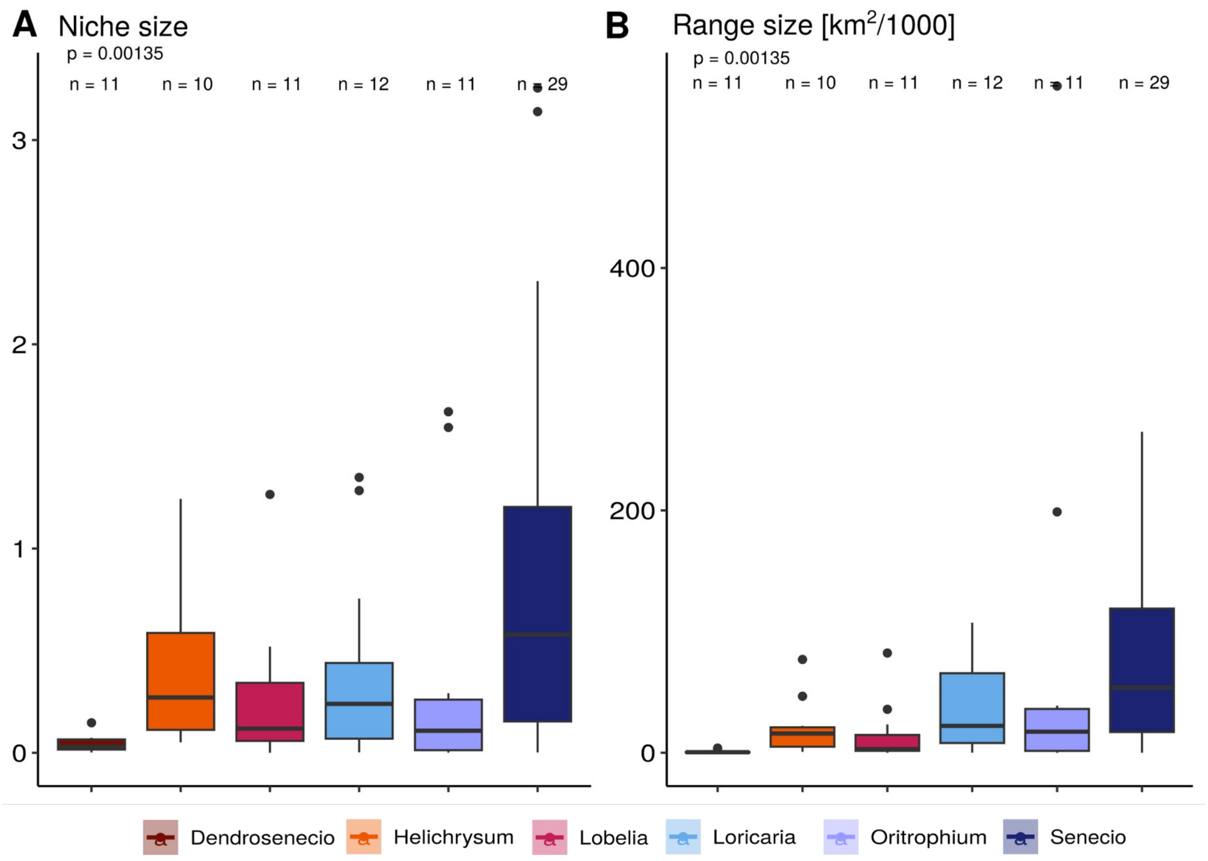

Figure S12. Niche and range size of tropical alpine lineages. (A) Boxplots of niche size and (B) range size summarised for all species per lineage. Boxplots show the median (centre line) and the first and third quartiles, while whiskers mark the 5th and 95th percentiles, values beyond these bounds are outliers. The p-values were calculated using Kruskal-Wallis test. Abbreviation: n - number of data points.

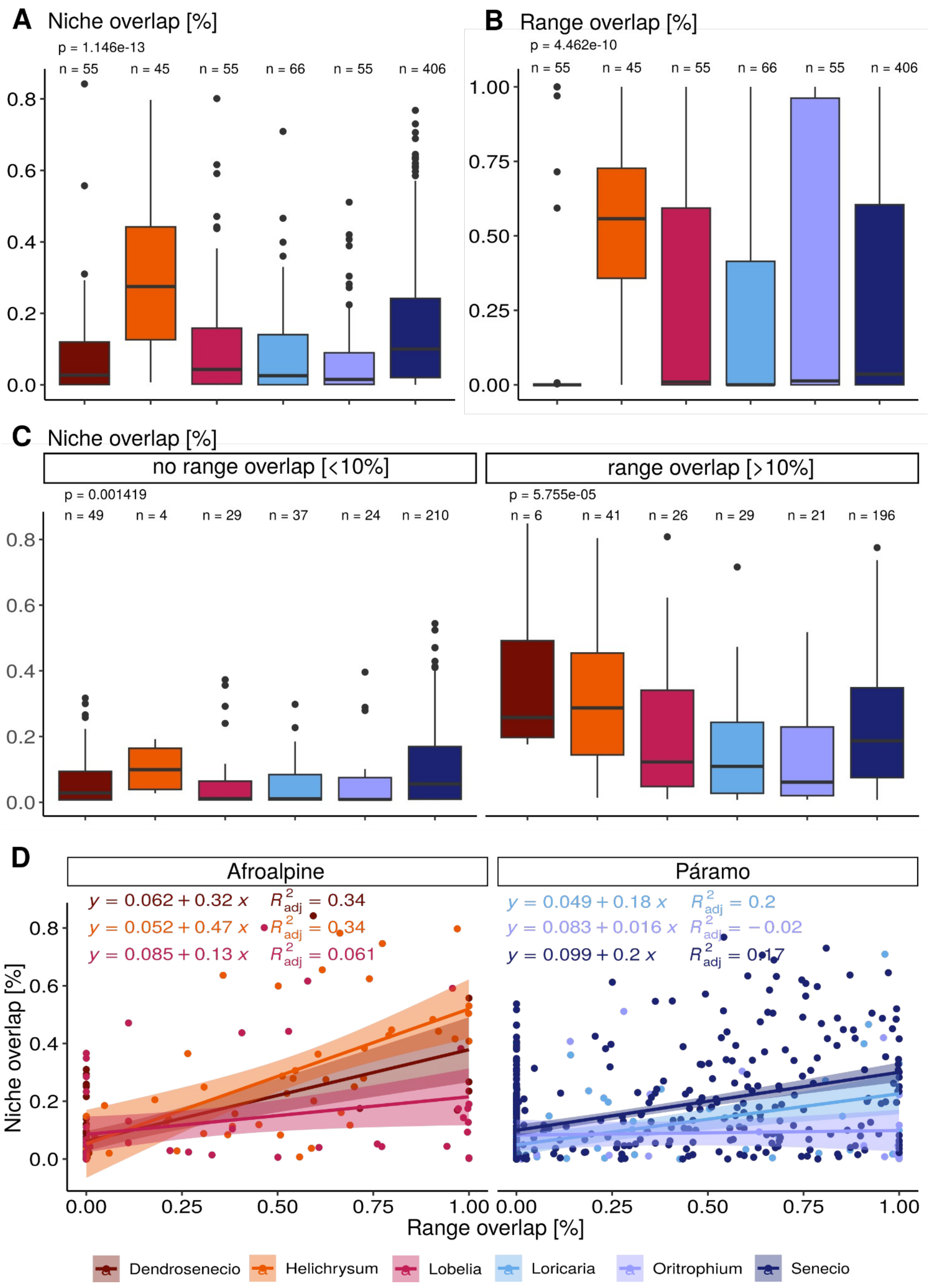

Figure S13. Niche and range overlap of tropical alpine lineages. (A) Boxplots of niche overlap and (B) range overlap summarised for all species-pairs per lineage. (C) Boxplot of niche overlap calculated for the subsets of species with non-overlapping and overlapping ranges. Boxplots show the median (centre line) and the first and third quartiles, while whiskers mark the 5th and 95th percentiles, values beyond these bounds are outliers. (D) Linear regression between range overlap and niche overlap separated per lineage. The p-values were calculated using Kruskal-Wallis test. Abbreviation: n - number of data points.

Table S6. Phylogenetic generalised least square analyses. Significance codes: 0 ‘***’ 0.001 ‘**’ 0.01 ‘*’ 0.05 ‘.’ 0.1 ‘ ’ 1. Abbreviation: DF - degree of freedom.

| | *Dendrosenecio* | | *Lobelia* | | *Helichrysum* | |
|---|---|---|---|---|---|---|
| Niche size ~ range size | intercept | slope | intercept | slope | intercept | slope |
| Estimate | 2.4999e-02 | 2.1746e-05 | 7.8341e-02 | 1.3077e-05 | 1.2916e-01 | 1.1068e-05 |
| Standard error | 1.1929e-02 | 7.6884e-06 | 6.3391e-02 | 2.2535e-06 | 1.2243e-01 | 3.8782e-06 |
| t-test | 2.0957 | 2.8284 | 1.2358 | 5.8032 | 1.055 | 2.854 |
| P-value | 0.06558 | 0.01977 | 0.2478009 | 0.0002584 | 0.32647 | 0.02455 |
| significant | . | * | | *** | | * |
| lambda median | 0.001 | | 0.001 | | 0.001 | |
| Residual standard error: | 0.0163 on 9 DF | | 0.07999 on 9 DF | | 0.1408 on 7 DF | |
| Multiple $R^2$/adjusted $R^2$ | 0.4706/ 0.4118 | | 0.7891/ 0.7657 | | 0.5378/ 0.4718 | |
| F-statistic/ p-value | 8 on 1 and 9 DF/ 0.01977 | | 33.68 on 1 and 9 DF/ 0.0002584 | | 8.145 on 1 and 7 DF/ 0.02455 | |
| Lilliefors normality test: D/ p-value | 0.278/ 0.01739 | | 0.21162/ 0.183 | | 0.16883/ 0.7201 | |

| | *Loricaria* | | *Oritrophium* | | *Senecio* | |
|---|---|---|---|---|---|---|
| Niche size ~ range size | intercept | slope | intercept | slope | intercept | slope |
| Estimate | 4.0580e-02 | 9.7165e-06 | 1.4891e-01 | 3.2594e-06 | -1.2732e-01 | 1.0063e-05 |
| Standard error | 1.5624e-01 | 2.7452e-06 | 1.2930e-01 | 6.5997e-07 | 2.2123e-01 | 1.2095e-06 |
| t-test | 0.2597 | 3.5394 | 1.1517 | 4.9387 | -0.5755 | 8.3200 |
| P-value | 0.801634 | 0.007626 | 0.287276 | 0.001678 | 0.571 | 4.366e-08 |
| significant | | ** | | ** | | *** |
| lambda median | 0.001 | | 0.001 | | 1.000 | |
| Residual standard error: | 0.1262 on 8 DF | | 0.1931 on 7 DF | | 0.0.2347 on 21 DF | |
| Multiple $R^2$/adjusted $R^2$ | 0.6103/ 0.5616 | | 0.777/ 0.7451 | | 0.7672/ 0.7562 | |
| F-statistic/ p-value | 12.53 on 1 and 8 DF/ 0.007626 | | 24.39 on 1 and 7 DF/ 0.001678 | | 69.22 on 1 and 21 DF/ 4.366e-08 | |
| Lilliefors normality test: D/ p-value | 0.18688/ 0.4136 | | 0.29824/ 0.02041 | | 0.14925/ 0.2032 | |

| Niche size ~ Niche size | *Dendrosenecio* | | *Lobelia* | | *Helichrysum* | |
|---|---|---|---|---|---|---|
| Axis1 | intercept | slope | intercept | slope | intercept | slope |
| Estimate | -0.024423 | 0.687078 | 0.041841 | 0.483656 | -0.098294 | 0.926882 |
| Standard error | 0.011956 | 0.099592 | 0.086777 | 0.114045 | 0.104292 | 0.172983 |
| t-test | -2.0428 | 6.8989 | 0.4822 | 4.2409 | -0.9425 | 5.3582 |
| P-value | 0.07144 | 7.076e-05 | 0.641194 | 0.002171 | 0.377324 | 0.001055 |
| significant | . | *** | | ** | | ** |
| lambda median | 0.356 | | 0.001 | | 0.001 | |
| Residual standard error: | 0.009413 on 9 DF | | 0.1006 on 9 DF | | 0.09168 on 7 DF | |
| Multiple $R^2$/adjusted $R^2$ | 0.841/0.8233 | | 0.6665/0.6294 | | 0.804/0.776 | |
| F-statistic/ p-value | 47.59 on 1 and 9 DF/ 7.076e-05 | | 17.99 on 1 and 9 DF/ 0.002171 | | 28.71 on 1 and 7 DF / 0.001055 | |
| Lilliefors normality test: D/ p-value | 0.21586/0.1621 | | 0.16942/0.5072 | | 0.11704/0.9761 | |

| Niche size ~ Niche size | *Loricaria* | | *Oritrophium* | | *Senecio* | |
|---|---|---|---|---|---|---|
| Axis1 | intercept | slope | intercept | slope | intercept | slope |
| Estimate | -0.28419 | 1.38658 | -0.19959 | 1.29880 | -0.251193 | 1.361598 |
| Standard error | 0.11033 | 0.12514 | 0.07248 | 0.10411 | 0.097466 | 0.090777 |
| t-test | -2.5759 | 11.0804 | -2.7537 | 12.4748 | -2.5772 | 14.9995 |
| P-value | 0.03282 | 3.927e-06 | 0.02835 | 4.899e-06 | 0.01757 | 1.077e-12 |
| significant | * | *** | * | *** | * | *** |
| lambda median | 1.000 | | 0.001 | | 0.001 | |
| Residual standard error: | 0.06534 on 8 DF | | 0.8484 on 7 DF | | 0.1476 on 21 DF | |
| Multiple $R^2$/adjusted $R^2$ | 0.9388/ 0.9312 | | 0.957/0.9508 | | 0.9146/0.9106 | |
| F-statistic/ p-value | 122.8 on 1 and 8 DF / 3.927e-06 | | 155.6 on 1 and 7 DF / 4.899e-06 | | 225 on 1 and 21 DF / 1.077e-12 | |
| Lilliefors normality test: D/ p-value | 0.16392/0.626 | | 0.23453/0.1639 | | 0.1351/0.3389 | |

| Niche size ~ Niche size | *Dendrosenecio* | | *Lobelia* | | *Helichrysum* | |
|---|---|---|---|---|---|---|
| Axis 2 | intercept | slope | intercept | slope | intercept | slope |
| Estimate | -0.027103 | 0.192840 | 0.21685 | 0.33638 | -0.19351 | 0.83126 |
| Standard error | 0.012993 | 0.031291 | 0.25779 | 0.29232 | 0.27106 | 0.37678 |
| t-test | -2.0859 | 6.1629 | 0.8412 | 1.1508 | -0.7139 | 2.2062 |
| P-value | 0.0666224 | 0.0001661 | 0.4220 | 0.2795 | 0.49838 | 0.06315 |
| significant | . | *** | | | | . |
| lambda median | 0.001 | | 0.816 | | 0.167 | |
| Residual standard error: | 0.009805 on 9 DF | | 0.1714 on 9 DF | | 0.1592 on 7 DF | |
| Multiple $R^2$/adjusted $R^2$ | 0.8084/0.7871 | | 0.1708/0.07863 | | 0.4102/0.3259 | |

| | | | | | |
|---|---|---|---|---|---|
| F-statistic/ p-value | 37.98 on 1 and 9 DF/ 0.0001661 | | 1.853 on 1 and 9 DF/ 0.2065 | | 4.867 on 1 and 7 DF / 0.06315 |
| Lilliefors normality test: D/ p-value | 0.17994/0.4091 | | 0.22537/0.1225 | | 0.16815/0.6548 |

| Niche size ~ Niche size Axis 2 | *Loricaria* | | *Oritrophium* | | *Senecio* | |
|---|---|---|---|---|---|---|
| | intercept | slope | intercept | slope | intercept | slope |
| Estimate | -0.29202 | 1.14318 | -0.34209 | 1.39518 | -0.12049 | 1.42343 |
| Standard error | 0.09635 | 0.12672 | 0.19017 | 0.27092 | 0.34701 | 0.29063 |
| t-test | -3.0308 | 9.0211 | -1.7989 | 5.1497 | -0.3472 | 4.8977 |
| P-value | 0.01629 | 1.821e-05 | 0.115066 | 0.001324 | 0.7319 | 7.635e-05 |
| significant | * | *** | | ** | | *** |
| lambda median | 0.001 | | 0.001 | | 1.000 | |
| Residual standard error: | 0.06047 on 8 DF | | 0.1869 on 7 DF | | 0.3323 on 21 DF | |
| Multiple $R^2$/adjusted $R^2$ | 0.9105/ 0.8993 | | 0.7912/0.7613 | | 0.5332/0.511 | |
| F-statistic/ p-value | 81.38on 1 and 8 DF / 1.821e-05 | | 26.52 on 1 and 7 DF / 0.001324 | | 23.99 on 1 and 21 DF / 7.635e-05 | |
| Lilliefors normality test: D/ p-value | 0.25716/0.0597 | | 0.2649/0.06832 | | 0.1576/0.1456 | |

*Supplementary Note S2. Phylogenetic reconstructions and molecular dating*

We selected six lineages from the families Asteraceae and Campanulaceae (all Asterales). Recently, phylogenies of all Asteraceae lineages based on Hyb-Seq data using the Compositae1061 loci[7] were generated[2,5,6,8,9]. For the Afroalpine clade of giant lobelias, we inferred a new phylogeny here using the same method as for the published phylogenies. For the methodology of sequencing and phylogenetic reconstruction please refer to [9]. Sampling information is provided in the Supplementary Tables S7 and S8. In the case of *Helichrysum,* we wanted to confirm monophyly of the species and exclude potential introgression events, thus we decided to calculate a phylogeny using multiple samples per tropical alpine species from the so-called “TA1 clade”. In the case of *Senecio*, we reused previously published data, but resequenced some low quality samples. For *Oritrophium,* we were provided with an unpublished phylogeny, which is an extension from [2], including six new species and a larger outgroup[10]; the methodology followed the same approach as above.

For all six lineages, we tested for introgression using Dsuite[11] following the approach in [9] and removed respective samples until no further introgression (>10%) was detected. We had to exclude three Páramo species from the final phylogenies because they showed signals of introgression. As a last step, we reduced all phylogenies to single samples per species, and removed taxonomically undescribed samples. For *Loricaria* and *Dendrosenecio*, the authors of the respective publications were so kind as to follow this approach and provide us with a phylogeny including one sample per species. Finally, ML phylogenies were used for dating (Table S10).

Table S7. Newly sampled material for this study and the corresponding voucher information. Details for different sequencing approaches (SA) are presented in Table S9. Herbarium codes according to Index Herbariorum: O - University of Oslo; BC - Institut Botànic de Barcelona; PRC - Charles University, Prague. Abbreviations: HC - Herbarium code; NA - not applicable; * - included as an outgroup sample; dupl. - duplicate.

| Genus | Species | Date collected | State | Locality | altitude [m] | Latitude | Longitude | Collector | SA | Seq Identifier | HC | Other labels | final sampling [0/1 - reason] |
|---|---|---|---|---|---|---|---|---|---|---|---|---|---|
| Dendrosenecio | battiscombei | 07/13/2009 | Kenya | Mt Kenya | 4009 | -0.1443 | 37.348917 | AFROALP II team | run5 | DE_050_5 | O | KN-1081-5 | 1 |
| Dendrosenecio | brassicifolius | 12/02/2009 | Kenya | Aberdare Mts: Mt Satima area. | 3865 | -0.31065 | 36.631917 | AFROALP II team | run5 | DE_006_2 | O | KN-0516-2 | 1 |
| Dendrosenecio | elgonensis | 29/01/2009 | Kenya | Mt Elgon: S of Mt Koitobos | 3629 | 1.100667 | 34.6215 | AFROALP II team | run5 | DE_003_4 | O | KN-0327-4 | 1 |
| Helichrysum | amblyphyllum | 20/01/2009 | Kenya | Mt Elgon: S of Mt Koitobos | 3915 | 1.105667 | 34.601833 | AFROALP II team | run5 | HE_041_2 | O | KN-0026_2/ O-DP-34828 | 0 - dupl. |
| Helichrysum | argyranthum | 04/09/2006 | Tanzania | from Nanokanoka village to the Olmoti crater | NA | -3.021178 | 35.679342 | M. Galbany & S. Arrabal | run5 | HE_016_x | BC | BC 867822/ BC 867822 | 0 - dupl. |
| Helichrysum | brownei | 11/09/2006 | Kenya | Mt Kenya, between Old Mosses Camp & Shipton's camp | NA | -0.082922 | 37.286269 | M. Galbany & S. Arrabal | run5 | HE_015_x | BC | BC 867843/ BC 867843 | 1 |
| Helichrysum | chionoides | 11/09/2006 | Kenya | Mt Kenya, between Old Mosses Camp & Shipton's camp | NA | -0.058306 | 37.291603 | M. Galbany & S. Arrabal | run5 | HE_019_x | BC | BC 867833/ BC 867833 | 1 |
| Helichrysum | ellipticifolium | 23/01/1970 | Kenya | Mt Kenya, W slope, Naro Moru Track | 2700-3100 | 0~10';S | 37~11'-13'E | A. Bjornstad | run5 | HE_025_x | O | A. Bjornstad 382 | 1 |
| Helichrysum | formosissimum | 15/10/2019 | Ethiopia | Simen Mts., road from Dilibza towards Ras Dejen. | 4117 | 13.22128 | 38.37089 | Speciation Clock | run5 | HE_006_2 | O | SC2019-ES-518-2/ O-DP-77520 | 1 |
| Helichrysum | stuhlmannii | 22/07/2019 | Uganda | Rwenzori Mts., Lower Bigo Bog. West of John Matte Hut. | 3435 | 0.38567 | 29.92527 | Speciation Clock | run5 | HE_061_1 | O | UR-409_1/ O-DP-74761-74770 | 0 - dupl. |
| Helichrysum | nandense | 09/02/1972 | Rwanda | Rwanda, road to de Rambura, env. du lac Kagogo, Gisenyi | 2300 | NA | NA | G. Troupin | runBC | BR15188892 | | G. Troupin 14373 | 0 - dupl. |
| Lobelia | giberroa | 25/10/2019 | Ethiopia | Simen Mts., from Debark on the way to Lib Bey, Lemalimo road. | 2631 | 13.19173 | 37.88357 | Speciation Clock | run3 | LO_005_5 | O | SC2019_ES-571_5 | 0 - dupl. |
| Lobelia | rhynchopetalum | 15/10/2019 | Ethiopia | Simen Mts., road from Dilibza towards Ras Dejen. | 4150 | 13.22294 | 38.36737 | Speciation Clock | run3 | LO_004_7 | O | SC2019_ES-507_7/ O-DP-77329 | 0 - dupl. |
| Lobelia | bambuseti | 16/02/2009 | Kenya | Aberdare Mts: along car road towards Satima. | 3466 | -0.339 | 36.668 | AFROALP II team | run6 | LO_019_1 | O | KN-0697_1/ O-DP-28398 | 1 |
| Lobelia | deckenii subsp. burtii | 28/11/2008 | Tanzania | Mt Meru: Saddle Hut area. | 3637 | -3.218 | 36.766833 | AFROALP II team | run6 | LO_029_1 | O | TZ-0408_1/ O-DP-38624 | 0 - dupl. |
| Lobelia | columnaris | 01/01/2007 | Cameroon | North-West Region, Bamenda Highlands. | NA | NA | NA | NA | run6 | LO_006_1 | O | AFR-603_1/ O-DP-63019 | 1 |
| Lobelia | deckenii subsp. deckenii | 22/11/2019 | Kenya | Mount Elgon, path from "Road end" to Koitobos Peak. | 3935 | 1.11595 | 34.60115 | Speciation Clock | run6 | LO_003_1 | O | SC2019_KE-795_1/ O-DP-78924 | 1 |
| Lobelia | deckenii | 12/02/ | Kenya | Aberdare Mts: at the end of the car road | 3588 | -0.30616 | 36.626 | AFROALP II team | run6 | LO_018_1 | O | KN-0525_1/ O- | 0 - |

| Genus | Species | Date collected | State | Locality | altitude [m] | Latitude | Longitude | Collector | SA | Seq Identifier | HC | Other labels | final sampling [0/1 - reason] |
|---|---|---|---|---|---|---|---|---|---|---|---|---|---|
| | subsp. gregoriana | 2009 | | towards Satima. | | | | | | | | DP-27637 | introgression |
| Lobelia | mildbraedii | 29/07/2008 | Uganda | Virunga Mts: Mt Muhavura, Kabaragnuma Swamp. | 3058 | -1.367483 | 29.671317 | AFROALP II team | run6 | LO_032_1 | O | UG-2171_1/ O-DP-40200 | 1 (sister to ETH) |
| Lobelia | stuhlmannii | 22/07/2019 | Uganda | Rwenzori Mts., upper end of Lower Bigo Bog, just after the end of the boardwalk. | 3436 | 0.38626 | 29.92187 | Speciation Clock | run6 | LO_035_1 | O | UR-411_1/ O-DP-74781 | 1 |
| Lobelia | telekii | 22/11/2019 | Kenya | Mount Elgon, path up to Koitobos Peak. | 4143 | 1.12366 | 34.60037 | Speciation Clock | run6 | LO_002_1 | O | SC2019_KE-787_1/ O-DP-78802 | 1 |
| Lobelia | aberdarica | 11/02/2009 | Kenya | Aberdare Mts: Mt Kinangop area. | NA | NA | NA | AFROALP II team | run6 | LO_017_1 | O | KN-0474_1/ O-DP-27431 | 1 |
| Lobelia | aberdarica | 19/11/2019 | Kenya | Mount Elgon, at Mutamaiu campsite. | 2814 | 1.06249 | 34.71824 | Speciation Clock | run7 | LO_001_9 | O | SC2019_KE-762_9/ O-DP-78490 | 0 - dupl. |
| Lobelia | achrochilus | 29/10/2018 | Ethiopia | Oromia, Dinsho, along the road to Web valley. | NA | 7.082962 | 39.781883 | Speciation Clock | run7 | LO_010_1 | O | EB-143_1/ O-DP-70950 | 0 - dupl. |
| Lobelia | achrochilus | 17/10/2008 | Ethiopia | Bale Mts: Dinsho. | 3281 | 7.05815 | 39.7657 | AFROALP II team | run7 | LO_011_1 | O | ET-1503_1/ O-DP-34375 | 1 |
| Lobelia | bambuseti | 16/06/2019 | Kenya | Mount Kenya, Naro Moru Route, just east from the Meteorological Station Campground. | 3067 | -0.1705833333 | 37.21468333 | Speciation Clock | run7 | LO_012_1 | O | KK-237_1/ O-DP-72409 | 0 - dupl. |
| Lobelia | deckenii subsp. bequartii | 22/07/2019 | Uganda | Rwenzori Mts., Lower Bigo Bog. West of John Matte Hut. | 3435 | 0.38567 | 29.92527 | Speciation Clock | run7 | LO_034_1 | O | UR-406_1/ O-DP-74721 | 0 - dupl. |
| Lobelia | deckenii subsp. bequartii | 30/07/2019 | Uganda | Rwenzori Mts., from Butawu camp to Kachope lake on Kilembe trail. | 3850 | 0.33802 | 29.87579 | Speciation Clock | run7 | LO_037_1 | O | UR-456_1/ O-DP-75245 | 1 |
| Lobelia | deckenii subsp. burtii | 30/11/2008 | Tanzania | Mt Meru: Betw. Saddle Hut & Miriakamba Hut. | 3589 | -3.217833 | 36.770667 | AFROALP II team | run7 | LO_030_1 | O | TZ-0498_1 /O-DP-39094 | 0 - dupl. |
| Lobelia | deckenii subsp. deckenii | 21/06/2019 | Kenya | Mount Kenya, south of Shipton's campsite. | 4298 | -0.14319 | 37.31584 | Speciation Clock | run7 | LO_014_1 | O | KK-272_1/ O-DP-72910 | 0 - dupl. |
| Lobelia | deckenii subsp. deckenii | 29/01/2019 | Tanzania | Kilimanjaro, Shira plateau. | 3642 | -3.04503 | 37.24776 | Speciation Clock | run7 | LO_027_1 | O | TK-205_1/ O-DP-71816 | 0 - dupl. |
| Lobelia | deckenii subsp. deckenii | 03/11/2008 | Tanzania | Mt Kilimanjaro: Shira Plateau near Mt Simba. | 3636 | -3.03425 | 37.243 | AFROALP II team | run7 | LO_028_1 | O | TZ-0025_1/ O-DP-37016 | 0 - dupl. |
| Lobelia | giberroa | 04/02/2009 | Kenya | Cherangani Hills: Mt Kamalagon. | 3100 | 1.176967 | 35.518383 | AFROALP II team | run7 | LO_016_1 | O | KN-0451_1/ O-DP-43831 | 0 - introgres |

| Genus | Species | Date collected | State | Locality | altitude [m] | Latitude | Longitude | Collector | SA | Seq Identifier | HC | Other labels | final sampling [0/1 - reason] sion |
|---|---|---|---|---|---|---|---|---|---|---|---|---|---|
| Lobelia | lindblomii | 16/02/2009 | Kenya | Aberdare Mts: at the end of the car road towards Satima. | 3619 | -0.3335 | 36.643167 | AFROALP II team | run7 | LO_020_1 | O | KN-0717_1/ O-DP-28491 | 1* |
| Lobelia | mildbraedii | 03/08/2019 | Uganda | Kigezi district, Kanaba, Echyua forest, Kanaba swamp next to the road Kabale-Kisoro. | 2298 | -1.2553 | 29.79626 | Speciation Clock | run7 | LO_040_1 | O | UR-486_1/ O-DP-75625 | 1 |
| Lobelia | rhynchopetalum | 28/10/2018 | Ethiopia | Oromia, on the way to Rira from Sanetti camp site, Sanetti plateau. | 4094 | 6.849243 | 39.889069 | Speciation Clock | run7 | LO_009_1 | O | EB-120_1/ O-DP-70790 | 1 |
| Lobelia | stuhlmannii | 31/07/2019 | Uganda | Rwenzori Mts., along path over Freshfield pass – Guy Yeoman camp. | 3598 | 0.34911 | 29.9124 | Speciation Clock | run7 | LO_039_1 | O | UR-471_1/ O-DP-75458 | 0 - dupl. |
| Lobelia | telekii | 19/06/2019 | Kenya | Mount Kenya, Naro Moru route, at the base of Batien Peak. | 4306 | -0.16199 | 37.30095 | Speciation Clock | run7 | LO_013_1 | O | KK-256_1/ O-DP-72670 | 0 - dupl. |
| Lobelia | thuliniana | 27/06/2009 | Tanzania | Mafinga Highlands: Matanana Village. | 1851 | -8.390667 | 35.184667 | AFROALP II team | run7 | LO_031_1 | O | TZ-0714_1/ O-DP-39290 | 1 |
| Lobelia | wollastonii | 28/07/2008 | Uganda | Virunga Mts: Mt Muhavura. | 4139 | -1.382767 | 29.677833 | AFROALP II team | run7 | LO_033_1 | O | UG-2173_1/ O-DP-40210 | 0 - dupl. |
| Lobelia | wollastonii | 23/07/2019 | Uganda | Rwenzori Mts., west of East Bukurungu lake, along Bukurungu trail. | 3815 | 0.40292 | 29.9428 | Speciation Clock | run7 | LO_036_1 | O | UR-419_1/ O-DP-74856 | 1 |
| Lobelia | lindblomii | 16/02/2009 | Kenya | Aberdare Mts: at the end of the car road towards Satima. | 3619 | -0.3335 | 36.643167 | AFROALP II team | run7 | LO_020_3 | O | KN-0717_3/ O-DP-28493 | 0* - dupl. |
| Lobelia | giberroa | 10/02/2022 | Rwanda | Volcanoes National Park, Muhavura volcano. | 2833 | 01.37134º | 29.69725º | M Galbany-Casals, JA Calleja, M Kandziora & E Ndayishimiye | run8 | LO_041_A | BC | GC-2777 | 1 |
| Lobelia | mildbraedii | 08/02/2022 | Rwanda | Volcanoes National Park, Mount Gahinga. | 2993 | 01º23,552 | 29º38,454 | M. Galbany-Casals, J. A. Calleja & M. Kandziora | run8 | LO_042_A | BC | GC-2761 | 0 - dupl. |
| Lobelia | mildbraedii | 12/02/2022 | Rwanda | Volcanoes National Park, Karisimbi volcano, from the campsite to the summit. | 3732 | 01.49714º | 29.46016º | M Galbany-Casals, JA Calleja, M Kandziora & E Ndayishimiye | run8 | LO_043_A | BC | GC-2805 | 0 - dupl. |
| Diplostephium | rupestre | 16/10/2008 | Ecuador | Carchi; Volcan Chiles, rocks above the swampy area, ca 0.5 km N of the antennas, ca 1 km N of the pass with the road Tufino - Maldonado. | 4157 | 0.8062777778 | -77.94222222 | P Sklenar, E Rejzkova, F Kolar | run5 | DIx003xX | PRC | 11517 | 1* |
| Oritrophium | orizabense | 15/02/2007 | Mexico | Veracruz Llave; north slope of Pico Orizaba, 5 km NE of summit, 3.5 km SW of Ejido Jacal, 9.5 km SW of Escola. | 3542 | 19.085 | -97.2467 | Pruski, J.F. & Ortiz, R.d. | run2 | 4171xPruski | PRC | Pruski 4171 | 1 |
| Oritrophium | aciculifolium | 17/07/1990 | Peru | Cajamarca; Cutervo, Fortaleza de Chontacruz, San Andres. | 2400 | NA | NA | S Llatas Quiroz & H Suárez C. | run2 | 2848xQuiroz | PRC | Quiroz 2848 | 1 |
| Oritrophium | yacuriense | 16/06/2019 | Ecuador | Loja; Cordillera las Lagunillas (de Sabanilla), paramo de las Lagunas Neg. | 3350 | -4.7091666 | -79.434166 | P Sklenar, J Mackova & P Macek | run2 | 12032 | PRC | 12032 | 1 |

| Genus | Species | Date collected | State | Locality | altitude [m] | Latitude | Longitude | Collector | SA | Seq Identifier | HC | Other labels | final sampling [0/1 - reason] |
|---|---|---|---|---|---|---|---|---|---|---|---|---|---|
| | | | | | | 67 | 67 | | | | | | |
| Oritrophium | repens | 12/11/2000 | Peru | | 3640 | -6.38 | -79.22 | Sanchez Vega | run3 | ORx005xx | PRC | Sanchez Vega 10342 | 1 |
| Oritrophium | ollgaardii | 01/28/2020 | Ecuador | Chimborazo; northern ridge of Altar. | 4350 | -1.637861111 | -78.43019444 | P. Sklenar | run3 | ORx014xx | PRC | 16079 | 1 |
| Oritrophium | llanganatense | 12/03/2010 | Ecuador | Tungurahua; Parque Nacional Llanganatis, slopes above laguna at the western side of Cerro Hermoso. | 4090 | -1.227277778 | -78.29875 | P. Sklenar | run2 | 13118 | PRC | 13118 | 1 |
| Oritrophium | peruvianum | 25/10/2008 | Ecuador | Chimborazo; Páramos around Cerro Quilimas, along the trail Alao - Huamboya, ca 3 km S of the bridge across Rio Alao. | 3492 | -1.83 | -78.45 | P Sklenar, E Rejzkova, F Kolar | run3 | ORx012xx | PRC | 11602 | 1 |
| Oritrophium | limnophilum | 25/10/2008 | Ecuador | Chimborazo; Páramos around Cerro Quilimas, along the trail Alao - Huamboya, ca 3 km S of the bridge across Rio Alao. | 3492 | -1.83 | -78.45 | P Sklenar, E Rejzkova, F Kolar | run3 | ORx009xx | PRC | 11603 | 1 |
| Senecio | rhizomatus | 11/01/2018 | Bolivia | Road La Paz-Coroico, about 3 km NE from the Cumbre pass, paramo Yungueno | 4320 | -16.315 | -68.02527778 | Sklenar P, Wojtasiak SS, Urfus T, Bartosova R | run3 | SE_015_x | | 15020 | 1 |

Table S8. Previously published sequences used in this study. Asterisk indicates samples used as outgroup.

| Phylogeny of | Genus | Species epithet | Sample identifier | Additional identifier | BioProject | final sampling [0/1 - reason] |
|---|---|---|---|---|---|---|
| Dendrosenecio | Cineraria | mazoensis | CI001xXH | | SAMN19185797 | 1* |
| | Emilia | abyssinica | EM001xXH | | SAMN19185798 | 1* |
| | Senecio | syringifolius | SS001xXH | | SAMN19185799 | 1* |
| | Adenostyles | alliariae | AD001xX | | SAMN19185796 | 1* |
| | Dendrosenecio | johnstonii | DE021x4 | | SAMN19185826 | 1 |
| | Dendrosenecio | kilimanjari | DE018x7 | | SAMN19185838 | 1 |
| | Dendrosenecio | meruensis | DE029x3 | | SAMN19185839 | 1 |
| | Dendrosenecio | keniensis | DE017x7 | | SAMN19185829 | 1 |
| | Dendrosenecio | keniodendron | DE013x10 | | SAMN19185831 | 1 |
| | Dendrosenecio | adnivalis | DE034x3 | | SAMN19185802 | 1 |
| | Dendrosenecio | ericirosenii | DE031x3 | | SAMN19185825 | 1 |
| | Dendrosenecio | cheranganiensis | DE004x5 | | SAMN19185811 | 1 |
| Oritrophium | Chiliotrichum | diffusum | 4000 | | SAMN11585365 | 1* |
| | Chrysocoma | ciliata | 94-37 | | SAMN11585366 | 1* |
| | Boltonia | diffusa | 5381 | | SAMN11585364 | 1* |
| | Conyza | sumatrensis | 12201 | | SAMN11585368 | 1* |
| | Erigeron | glaucus | 447 | | SAMN11585371 | 1* |
| | Oritrophium | hieracioides | 15024 | | SAMN28597246 | 1 |
| | Oritrophium | ferrugineum | 738xKunkel | | SAMN28597245 | 1 |
| | Oritrophium | mucidum | 12214 | | SAMN28597248 | 1 |
| | Oritrophium | crocifolium | ORx015xA | | SAMN28597244 | 1 |
| Loricaria | Gnaphalium | antennarioides | 12412 | | SAMN22857824 | 1* |
| | Antennaria | anaphaloides | 134858x18xS22 | | SAMN11585394 | 1* |
| | Cuatrecasasiella | issernii | 12405 | | SAMN22857823 | 1* |
| | Luciliocline | subspicata | 3xS3 | | SAMN11585410 | 1* |
| | Loricaria | graveolens | LRx017xXH | | SAMN22857848 | 1 |
| | Loricaria | unduaviensis | LRx001xx | | SAMN22857874 | 1 |
| | Loricaria | lycopodinea | LRx020xXH | | SAMN22857855 | 1 |
| | Loricaria | ferruginea | LRx005xXH | | SAMN22857845 | 1 |
| | Loricaria | colombiana | LRx008xX | | SAMN22857838 | 1 |
| | Loricaria | puracensis | LRx006xB | | SAMN22857836 | 1 |
| | Loricaria | ilinissae | 11563 | | SAMN22857851 | 1 |
| | Loricaria | antisanensis | LRx007xC | | SAMN22857835 | 1 |
| | Loricaria | pauciflora | LRx009xX | | SAMN22857856 | 1 |
| | Loricaria | thuyoides | 11529 | | SAMN22857872 | 1 |
| | Loricaria | scolopendra | 11610 | | SAMN22857859 | 1 |
| | Loricaria | complanata | 14281xBetancour | | SAMN22857842 | 1 |
| Helichrysum | Helichrysum | amblyphyllum | HE_003_1 | SC2019-KE-784-1 | SAMN33382454 | 1 |
| | Helichrysum | argyranthum | HE_017_x | BC 867828 | SAMN33382463 | 1 |
| | Helichrysum | brownei | HE_047_1 | KN-911_1 | SAMN33382482 | 0 - dupl. |
| | Helichrysum | chionoides | HE_036_x | BC 867840 | SAMN33382494 | 0 - dupl. |
| | Helichrysum | ellipticifolium | HE_043_2 | KN-0483-2 | SAMN33382529 | 0 - dupl. |
| | Helichrysum | formosissimum | HE_005_1 | SC2019-KE-767-1 | SAMN33382542 | 0 - dupl. |
| | Helichrysum | mariepscopicum | HE_009_x | R-14592 | SAMN33382612 | 1* |
| | Helichrysum | meyeri-johannis | HE_020_x | BC 867837 | SAMN33382619 | 1 |
| | Helichrysum | nandense | HE_023_x | Lye 1492 | SAMN33382639 | 1 |
| | Helichrysum | newii | HE_001_1 | SC2019-TK-172-1 | SAMN33382643 | 1 |
| | Helichrysum | reflexum | HE_010_x | R-14571 | SAMN33382681 | 1* |
| | Helichrysum | stuhlmanii | HE_022_x | BC 867841 | SAMN33382714 | 1 |
| | Helichrysum | elegantissimum | R14435 | | SAMN33382527 | 1* |
| | Helichrysum | flammeiceps | K1854 | | SAMN33382538 | 1* |
| | Helichrysum | foetidum | R14464 | | SAMN33382541 | 1* |
| | Helichrysum | heterolasium | RJBSAF03018 | | SAMN33382571 | 1* |
| | Helichrysum | kirkii | BR9615656 | | SAMN33382586 | 1* |
| | Helichrysum | korongoni | AS19 | | SAMN33382587 | 1* |
| | Helichrysum | patulifolium | BR14603303 | | SAMN33382662 | 1* |
| | Helichrysum | ruandense | BR14604430 | | SAMN33382687 | 1* |
| *Senecio* ser. *Culcitium* | Senecio | culcitioides | 11512 | | SAMN41274980 | 1 |
| | Senecio | comosus | SE_051_H | B10843218 | SAMN41274976 | 1 |
| | Senecio | algens | SE_050_X | 15091 | SAMN41274960 | 1 |
| | Senecio | candolleii | 1584xZuloaga | | SAMN41274968 | 1 |
| | Senecio | pflanzii | SE_012_x | Beck 9094 | SAMN41275036 | 1 |
| | Senecio | expansus | 16304xZuloaga | | SAMN41274986 | 0 - introgression |

| Phylogeny of | Genus | Species epithet | Sample identifier | Additional identifier | BioProject | final sampling [0/1 - reason] |
|---|---|---|---|---|---|---|
| | Senecio | hyoseridis | SE_014_x | Lewis 2579 | SAMN41274993 | 1 |
| | Senecio | praeruptorum | SE_013_x | Tupayachi 644125 | SAMN41275039 | 1 |
| | Senecio | tephrosioides | SE_018_x | SanchezVega 11401 | SAMN41275058 | 1 |
| | Senecio | serratifolius | SE_055_xH | B10745148 | SAMN41275047 | 1 |
| | Senecio | canescens | 15259 | | SAMN41274970 | 1 |
| | Senecio | violifolius | SE_056_xH | B10745138 | SAMN41275060 | 1 |
| | Senecio | campanulatus | SE_019_x | Saldias 4620 | SAMN41274964 | 1 |
| | Senecio | pindilicensis | SE_032_x | 11115 | SAMN41275038 | 1 |
| | Senecio | betonicifolius | SE_026_A | 12024 | SAMN41274982 | 1 |
| | Senecio | nivalis | SE_044_x | 16053 | SAMN41275026 | 1 |
| | Senecio | mojandensis | SE_030_A | 11028 | SAMN41275023 | 1 |
| | Senecio | longepenicillatus | 10195 | | SAMN41275016 | 0 - introgression |
| | Senecio | eliseae | SE_042_x | 15500 | SAMN41274985 | 1 |
| | Senecio | ferrugineus | 9348 | | SAMN41274988 | 1 |
| | Senecio | cocuyanus | S3_A | 12212 | SAMN41274975 | 1 |
| | Senecio | hypsobates | SE_007_x | Idobro 3160 | SAMN41274995 | 1 |
| | Senecio | otophorus | 12267 | | SAMN41275031 | 1 |
| | Senecio | cuencanus | SE_025_A | 11128 | SAMN41274979 | 1 |
| | Senecio | josei | SE_043_x | 15701 | SAMN41275008 | 1 |
| | Senecio | superandinus (quitensis) | SE_035_A | 11509 | SAMN41275041 | 0 - dupl. |
| | Senecio | lingulatus | SE_033_x | 11162 | SAMN41275015 | 0 - introgression |
| | Senecio | imbaburensis | SE_008_x | Cazalet 5775 | SAMN41274997 | 1 |
| | Senecio | gargantanus | SE_028_A | 54 | SAMN41274990 | 1 |
| | Senecio | puracensis | S19 | 12311 | SAMN41275040 | 0 - introgression |
| | Senecio | superparamensis | SE_027_A | 11544 | SAMN41275057 | 0 - introgression |
| | Senecio | keshua | SE_010_x | Teiller 6471 | SAMN41275011 | 1 |
| | Senecio | patens | SE_031_B | 11113B | SAMN41275034 | 0 - introgression |
| | Senecio | antisanae | S10 | 12094 | SAMN41275004 | 0 - dupl. |
| | Senecio | involucratus | SE_036_A | 11566 | SAMN41275001 | 1 |
| | Senecio | antisanae | SE_001_x | Palacios 7384 | SAMN41274961 | 1 |
| | Senecio | superandinus | SE_034_A | 11188 | SAMN41275055 | 1 |
| | Senecio | magellanicus | SEx011xx | Zuloaga 14064 | SAMN41275019 | 1* |
| | Senecio | gilliesii | SEx006xx | Salariato 145 | SAMN41274992 | 1* |
| | Senecio | candidans | 1079xMezaMartinez | | SAMN41274965 | 1* |
| | Senecio | martinensis | 189xRatto | | SAMN41275020 | 1* |
| | Robinsonia | gayana | ROx004xXH | 11985 | SAMN41274957 | 1* |

Table S9. Information on sequencing runs, number of samples during sequencing, sequencing maschine, and library preparation adjustments. Abbreviations: NA - not available.

| run | number of samples | Genera sequenced | sequencer | spiked in unenriched libraries |
|---|---|---|---|---|
| runBC | NA | Helichrysum | NovaSeq | 40% |
| run1 | 24 | Senecio | MiSeq | 0% |
| run2 | 96 | Loricaria, Oritrophium | NextSeq | 30% |
| run3 | 96 | All studied genera | NovaSeq | 50% |
| run5 | 88 | All studied genera | NovaSeq | 40% |
| run6 | 94 | Lobelia and others | NovaSeq | 40% |
| run7 | 20 | Lobelia | MiSeq | 30% |
| run 8 | 80 | All studied genera | NovaSeq | 40% |

Table S10. Age estimates in millions of years (Ma) used as secondary calibration points for the molecular dating.

| lineage | Calibration in Ma | Notes on calibration points |
|---|---|---|
| *Helichrysum* | TA1 stem: 9.8 - 10.2[6] | No confidence interval provided, adding a |
| | H. chinoides-H. amblyphyllum: 3.5-3.9[6] | range of 0.2 Ma for min and max calibrations. |
| *Lobelia* | Crown Giant: 15.17 - 5.00[12] | Crown Giants: based on BEAST result. |
| | Root: 9.63-22.98[12] | Root: stem age of clade that includes giant lobelias based on BEAST result.. |

| | | |
|---|---|---|
| *Dendrosenecio* | Root: 33.55-19.75[12] | Based on BEAST result. |
| | Dendrosenecio crown: 16.77 - 2.07[12] | |
| *Loricaria* | Root: 12.0 - 6.0[13] | Chosen as node between 26 and 25. |
| *Oritrophium* | Root: max 30.0[14] | |
| | *Oritrophium* stem: 21.0 - 3.75[15] | |
| *Senecio* | Root: 9 - 3.1[16] | |

*Supplementary Note S3. Occurrence and niche information*

The occurrence information for the Páramo lineages were recorded from the following sources over the course of the year 2021: Tropicos, Botanical Collection from the Field Museum, The Kew Herbarium Catalogue, Botany Collections from the Smithsonian National Museum of Natural History, Herbario Virtual de la Universidad Nacional de Colombia, and Instituto de Botánica Darwinion. Additionally, species occurrences were obtained from The Flora and Vegetation Database for Andean Páramo[17] and from the personal database of one of the co-authors, Petr Sklenář, which includes his botanical collections (mostly deposited at the Herbarium at Charles University in Prague, PRC) and photographs of herbarium vouchers taken at different herbaria in Venezuela and at the Herbarium of Universidad Nacional in Colombia. The raw data was meticulously curated. If the photograph of the herbarium voucher was available, the species identity was verified. If no photograph of the collection was available and the identity of the collector or the person who determined the species could not be verified, the database entry was discarded. For endemic species with restricted distribution, we opted to construct the geographic coordinates based on the geographic information provided directly on the voucher. For Páramo taxa with insufficient occurrence information, we added occurrences from GBIF, querying each species name in late 2022. For Afroalpine species, we mainly used the collections from the collection at the herbarium of the University of Oslo (O), and supplemented them with GBIF data in Summer 2022 by querying species names. During the filtering of occurrence records we did not reduce the data to one occurrence point per grid, as this would have led to an enormous reduction for narrowly endemic species. Finally, each occurrence point was projected onto a map for validation and removed if it was far outside the known range of the species. All lineages are well-known representatives of the tropical alpine ecosystem. Most species of the South American lineages are restricted to the Páramo, but some species from adjacent areas were included in our analysis to ensure monophyly.

To calculate the niche metrics, we first reduced the 19 bioclimatic variables from CHELSA[18] to minimise correlation using the vifstep function of the R package usdm v.2.1-7[19], aiming for a VIF below 10. Bioclimatic variables were transformed beforehand using the natural logarithm to scale among variables. Species occurrences were projected onto the $PCA_{env}$ to obtain PC scores that are spatially and phylogenetically unbiased[20,21].

**Supplementary Data S1. associated digital data**

See zip file/folder
